# Healthcare Infrastructure Shapes Evolutionary Trade-offs and Geographic Dissemination of Multidrug-Resistant *Acinetobacter baumannii*

**DOI:** 10.64898/2026.01.06.697983

**Authors:** Shengkai Li, Yilei Wu, Yan Zhou, Ling Zhong, Yongmei Jiang, Yuezhuo Wang, Jin Li, Haixia Lin, Heng Li, Shuting Xia, Hong Du, Rong Zhang, Yongliang Lou, Shaopu Wang, Acinetobacter baumannii Research Group, Ping He, Min Wang, Jimei Du, Zhemin Zhou

**Affiliations:** MOE Key Laboratory of Geriatric Diseases and Immunology, Department of Clinical Laboratory, Second Affiliated Hospital of Soochow University, Cancer Institute, Suzhou Medical College, Soochow University, Suzhou 215127, China; National Center of Technology Innovation for Biopharmaceuticals, Suzhou Biomedical Industry Innovation Center, Suzhou 215127, China; Department of Life Sciences, Imperial College London, London SW7 2AZ, UK; Department of Microbiology and Immunology, School of Laboratory Medicine and Life Science, Institute of One Health, Wenzhou Key Laboratory of Sanitary Microbiology, Wenzhou Medical University, Wenzhou 325035, China; Department of Laboratory Medicine, West China Second University Hospital, Sichuan University, Chengdu 610000, China; Key Laboratory of Birth Defects and Related Diseases of Women and Children (Sichuan University), Ministry of Education, West China Second University Hospital, Sichuan University, Chengdu 610000, China; Department of Clinical Laboratory, Second Affiliated Hospital of Zhejiang University, School of Medicine, Hangzhou 310000, China; Department of Pediatrics, West China Second University Hospital, Sichuan University, Chengdu 610000, China; Department of Immunology and Microbiology, School of Medicine, Shanghai Jiao Tong University, Shanghai 201100, China; National Key Laboratory of Intelligent Tracking and Forecasting for Infectious Diseases, National Institute for Communicable Disease Control and Prevention, Chinese Center for Disease Control and Prevention, Beijing 102207, China

## Abstract

Antimicrobial-resistant pathogens pose an existential threat to modern medicine, yet the evolutionary forces driving their adaptation in healthcare systems remain largely unexplored. We revealed that hospital network architecture functions as a primary selective pressure, driving pathogen evolution through infrastructure-dependent virulence-transmission trade-offs. Phylogenomic analysis of 5,023 *Acinetobacter baumannii* isolates across China’s centralized healthcare system identifies two co-existing evolutionary strategies: a virulence-optimized clade (ESL2.4) that spread slowly (20.4 km per year) in low-connectivity hospitals, and a transmission-optimized clade (ESL2.5) that disseminate rapidly (65.2 km per year) through mega-city healthcare hubs, likely attributed to its capsule conversion and increased upper respiratory colonization. Comparative analysis with European *A. baumannii* populations demonstrates that healthcare connectivity, not geography, governs pathogen distribution through convergent genomic adaptations. Our simulation suggests competitive asymmetries of the two clades, following the ecotype principle and enabling stable coexistence. The COVID-19 pandemic provided a natural experiment validating these mechanisms: outpatient restrictions reduced transmission-optimized lineage spread by 89%, while virulence-optimized lineage persisted through inpatient networks. These findings establish healthcare infrastructure as a critical evolutionary driver with immediate implications for predicting and controlling antimicrobial resistance emergence across diverse healthcare systems.

## Introduction

The evolutionary transition of *Acinetobacter baumannii* from environmental saprophyte to formidable nosocomial pathogen exemplifies how healthcare environments drive rapid bacterial adaptation^1^. This transformation has been particularly pronounced in developing countries, where rapid expansion of hospital infrastructure has coincided with alarming rises in multidrug-resistant *A. baumannii* (MDR-AB) infections^2,3^. Yet how healthcare system architecture influences pathogen evolution and whether it generates predictable evolutionary trade-offs remains poorly understood^4,5^.

Classical evolutionary theory predicts that selection for between-host transmissions typically constrains within-host virulence, preventing pathogens simultaneously maximizing virulence and transmissibility^5,6^. While such virulence-transmission trade-off has been extensively documented across viral pathogens^7^, bacterial populations often evolve distinct ecotypes adapted to different niches^8^, contributing to the maintenance of bacterial metapopulations and eventual speciation^9^. Healthcare infrastructure likely imposes analogous selective pressures for antimicrobial resistant (AMR) bacterial pathogens, yet their roles in shaping evolutionary trade-offs and ecological diversification remain underexplored.

Global surveys have identified multiple co-circulating clades within international clone 2 (IC2), the predominant *A. baumannii* lineage^2^, but lack integration with healthcare-system data, precluding resolution of how these clades evolved from healthcare architecture. China provides an exceptional natural experiment: its large geographic scale, dense and heterogeneous healthcare network, and characteristic patient mobility patterns, including frequent self-referral to tertiary hospitals, generate a diversity of transmission environments that differ from referral-hierarchy systems in many high socio-economic countries^10,11^. This creates distinct selective pressures that may favor different combinations of virulence, persistence, and transmission capabilities across hospital types and geographic regions.

Here we integrated >5,000 genomes with comprehensive healthcare data to reveal how hospital infrastructure shapes *A. baumannii* evolution in China. Contemporary Chinese *A. baumannii* populations are shaped through two mechanisms: connectivity-dependent geographic distribution patterns coincide with human socio-economic interactions and differential selection for evolutionary strategies optimized for distinct transmission environments. Mega-city hubs select for rapid-spreading, persistent lineages optimized for upper respiratory tract colonization, while peripheral facilities favor virulent but geographically constrained strains. Comparative analysis with *Klebsiella pneumoniae* and European *A. baumannii* reveals convergent patterns, suggesting conserved adaptation strategies among hospital-acquired pathogens. Our findings underscore healthcare infrastructure as a primary driver of pathogen evolution, with immediate implications for surveillance and control strategies.

## Results

### Healthcare Architecture Shapes Pathogen Population Structure Through Connectivity-Dependent Selection

We assembled a comprehensive collection of 5,023 Chinese *Acinetobacter baumannii* genomes by sequencing 1,042 new isolates from 20 cities across 15 provinces and integrating them with 3,981 publicly available genomes, yielding broad geographic coverage across 73 cities in 27 provinces (1999–2024, **Table S1**). Phylogenetic analysis revealed overwhelming IC2 predominance in China (>90%, **Fig. 1a, Fig. S1**). Using SNP barcodes derived from previous study^2^, we resolved Chinese IC2 into three globally recognized clades of Epidemic Super Lineage (ESL) 2.5 (~50%), ESL2.4 (~48%), and ESL2.3 (~2%).

**Fig. 1:**
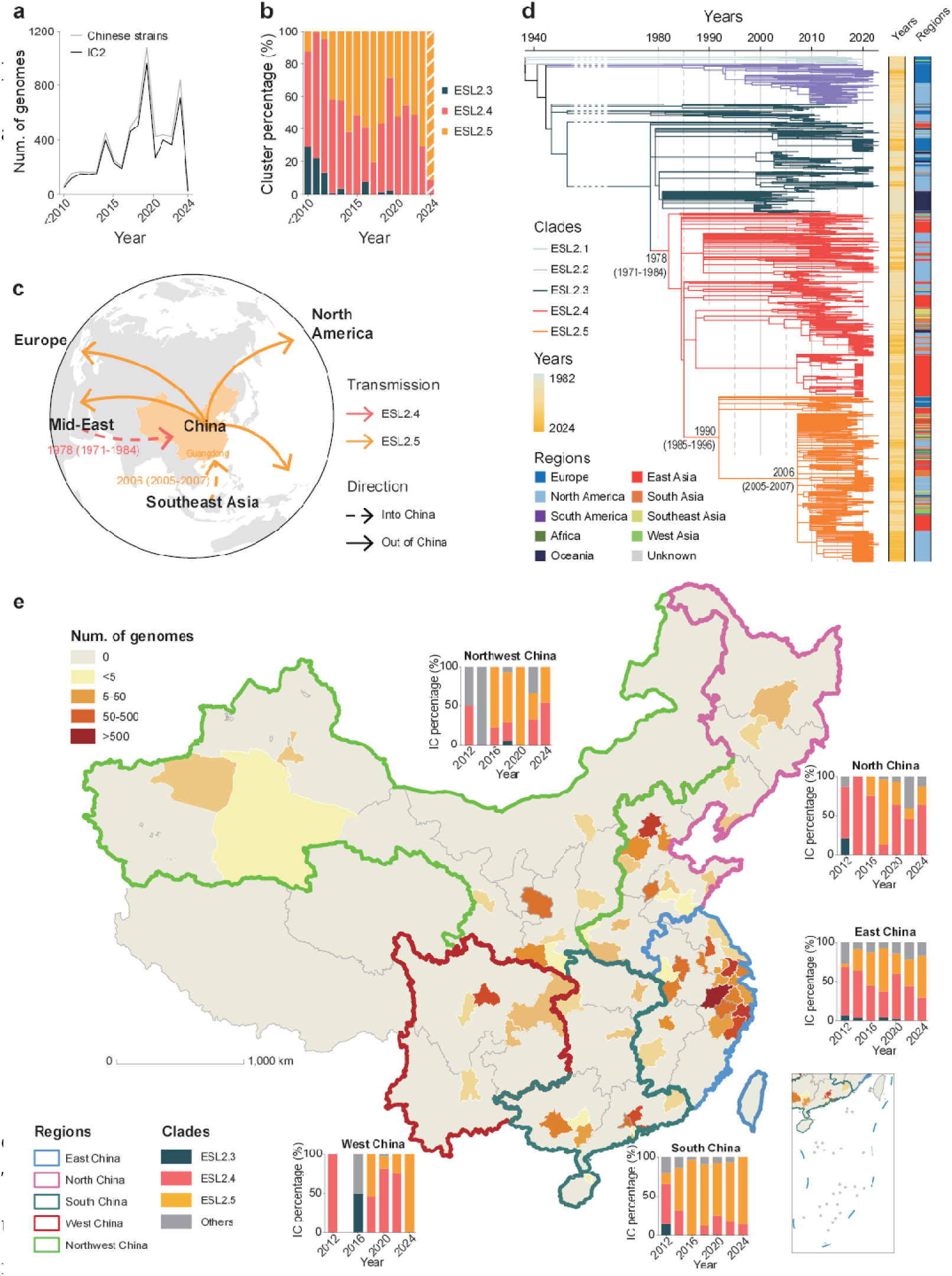
Temporal, phylogenetic and geographic dynamics of IC2 clades in China. **a,** Annual number of *A. baumannii* genomes included in this study from China (grey), overlaid with the number of IC2 genomes (black) sampled between 2010 and 2024. **b,** Temporal changes in the relative abundance of major IC2 clades in China. Stacked bars show the yearly proportions of ESL2.3 (teal), ESL2.4 (pink), and ESL2.5 (orange) among IC2 isolates. Hatched bars indicate years with limited sample sizes. **c,** Inferred interregional transmission routes of ESL2.4 and ESL2.5. Arrows indicate the predominant direction of lineage movement between regions of ESL2.4 (red) and ESL2.5 (orange). Approximate introduction periods are indicated for major transmission events. **d,** Time-calibrated phylogeny of global IC2 isolates. Branches are coloured by clade assignment. Right-hand annotations indicate sampling year (left bar) and geographic region (right bar) for each isolate. **e,** Spatial distribution and regional dynamics of IC2 clades across China. Provinces are coloured according to the number of genomes sampled. Temporal compositions of IC2 clades in each major geographic region (Northwest, North, East, South, and West China) from 2012 to 2024 are shown as stacked bars. Provincial boundaries and regional groupings are indicated. Scale bar is shown in kilometres (km).

Temporal reconstruction demonstrates a marked shift in clade composition over the last two decades. ESL2.4 was the dominant lineage in China during the early 2010s; however, ESL2.5 increased rapidly from ~ 10% to 78% by 2017, followed by stabilization with prevalence fluctuating around 60% thereafter (**Fig. 1b**). Analyses of contemporaneous genomes from Europe (1229 genomes) and the United States (5681 genomes) reveal parallel expansion-and-stabilization patterns for ESL2.5 over 10–15 years, indicating a global trend. Phylogeographic inference estimates China as a key hub, as the major ESL2.5 subgroup (ESL2.5.6) reached Guangdong soon after its inferred emergence in Southeast Asia (circa 2004–2006) and subsequently dispersed into over 41 countries globally (**Fig. 1c**).

### Temporal Dynamics Reveal Distinct Evolutionary Trajectories and Introduction Events

Phylogenetic reconstruction reveals contrasting spatiotemporal histories for the two dominant clades. ESL2.4 appears as a long-established, locally endemic lineage with roots traceable to the 1970s–1980s, consistent with decades of persistence. In contrast, ESL2.5 emerged more recently in the late 1990s to early 2000s and underwent rapid geographic expansion (**Fig. 1c, d**). After initial establishment in coastal regions (2006–2008), ESL2.5.6 spread to central provinces by 2012 and achieved near-complete nationwide distribution by 2015. Across all major economic regions (East, North, South, West, Northwest), ESL2.4 initially dominated but was progressively out-competed by ESL2.5 before 2020 (**Fig. 1b, e**).

Despite its epidemiological dominance, ESL2.5 shows no appreciable AMR advantage over ESL2.4. Genotypic and phenotypic profiling revealed comparable resistance to all tested antibiotics (**Fig. S2**), with only subtle differences in PBP3 mutations and an ADC β-lactamase variant^2^. This near-equivalence in AMR determinants indicates that resistance mechanisms alone cannot explain ESL2.5’s rapid rise, implicating other evolutionary drivers.

### Geographic Transmission Networks Reveal Hub-Dependent Dissemination Patterns

We reconstructed *A. baumannii* transmission networks across China without imposing prior assumptions. Five major transmission module emerged, each spanning ~800 km and separated by sharply reduced inter-module connectivity (462 versus 151; **Fig, 2a, b** and **Fig. S3a**). These modules map closely onto China’s principal economic regions: East (Yangtze River Delta), South (Pearl River Delta), Southwest (Chengdu Basin), North (Beijing–Tianjin), and Northwest. East China exhibited the densest internal networks, reflecting how urbanization and healthcare concentration jointly shape high-connectivity transmission corridors (**Fig. S4**).

**Fig. 2:**
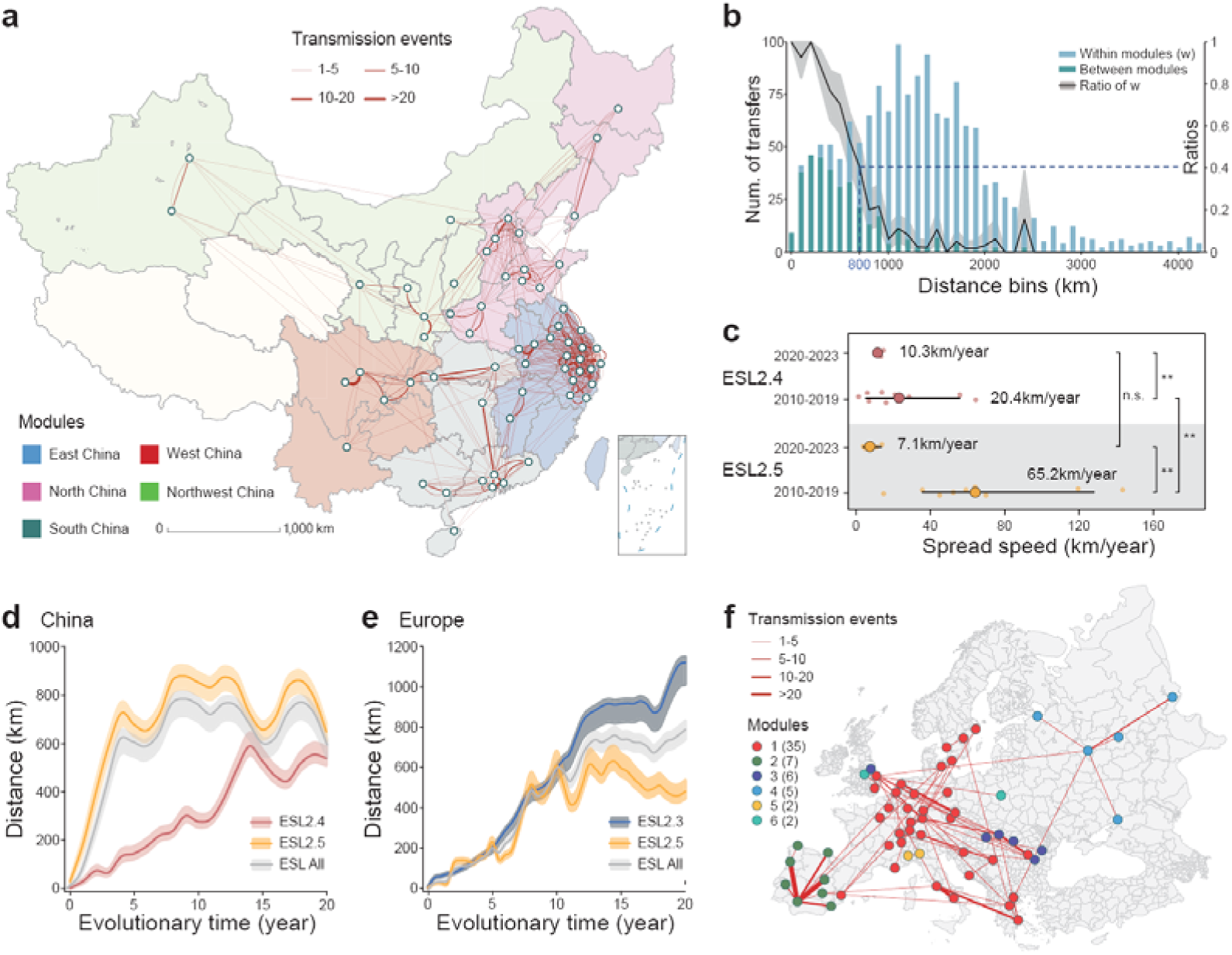
Hub-dependent transmission architecture and dissemination dynamics. **a,** Reconstructed transmission network of *A. baumannii* across China. Nodes represent cities and edges represent inferred transmission events between cities. Edge thickness indicates the number of transmission events. Colours of provinces denote transmission modules corresponding to major geographic regions (East, North, South, West, and Northwest China). Scale bar is shown in kilometres (km). **b,** Distance-dependent structure of transmission events. Bars show the number of inferred transmission events occurring within modules (dark blue) or between modules (light blue) across geographic distance bins. The black line indicates the ratio of within-module events (*w*), with shaded areas representing 95% confidence intervals derived from bootstrap resampling. The red dashed line marks the distance (~800 km) at which within- and between-module events occur with equal probability. **c,** Estimated dissemination speeds of ESL2.4 and ESL2.5 in China inferred from phylogenetic branch lengths and geographic displacement. Horizontal bars indicate median values. Comparisons between time periods and clades were assessed using non-parametric tests (\*\**P* < 0.01; n.s. not significant). **d-e,** Accumulated geographic spread distance of IC2 clades as a function of evolutionary time in China (**d**) and Europe (**e**). Curves represent mean distances for ESL2.3 (blue), ESL2.4 (red), ESL2.5 (orange), and all IC2 isolates combined (grey). Shaded bands indicate 95% confidence intervals. **f,** Transmission network of *A. baumannii* in Europe, revealing a less centralized and more inter-connected network structure compared with China. An interactive, dynamic version of the transmission dynamics is available online at https://naclist.github.io/naclist-portfolio/sources/ab_china_dynamics/.

Node degree distributions fit a power-law (R² = 0.85), consistent with scale-free architecture dominated by a small number of highly connected hubs (**Fig. S4b**). This topology accords with observed patient referral patterns to major tertiary centers^12^. City connectivity correlated strongly with both population size and health-care resource density, underscoring the central role of mega-city hospitals as dissemination hubs. Smaller municipalities preferentially linked to provincial capitals rather than to geographically closer peers (**Fig. S3f, Table S2**), mirroring health-seeking behaviors that bias flows toward tertiary facilities.

### *A. baumannii* Clades Exhibited Different Dissemination Rates

To quantify dissemination efficiency, we estimated spatial migration distance as a function of evolutionary time (**See Methods**). ESL2.5 exhibited consistently greater geographic expansion (~800 km in 10 years) than ESL2.4 (~500 km in 20 years; **Fig. 2d**). Annual migration velocities revealed a similar contrast. ESL2.5 reached a median speed of 65.2 km·yr_¹ (range: 20-150 km·yr_¹) in the 2010s, compared with 20.4 km·yr_¹ for ESL2.4 (range: 2–70 km·yr_¹; **Fig. 2c**). Importantly, both clades utilized the same hub-dependent transmission networks, differing in velocity rather than route architecture. This implies clade-specific ecological or phenotypical differences that modulate spread efficiency within structurally constrained pathways.

### Healthcare System Architecture Determines Transmission Dynamics

To test whether these transmission signatures reflect health-system architecture rather than pathogen identity, we performed comparative phylogeographic analysis using 2,800 publicly available *A. baumannii* genomes sampled from 12 European countries (2010–2023, **Table S1**).

The European dataset revealed contrasting architecture. Networks were more decentralized and more inter-connected over long distances (**Fig. 2f**). Geographic distance constrained transmission much less in Europe (distance–transmission correlation *r* = −0.18, *p* = 0.08) than in China (*r* = −0.67, *p* < 0.001), consistent with a system in which cross-border and interregional patient flows and referral patterns reduce the spatial decay of transmission.

We recovered four major transmission modules in Europe. Two modules (in Spain and East Europe) remained isolated, while West and Central European countries formed two highly entangled modules lacking clear geographical specificity. The mean clustering coefficient in Europe (0.31) was substantially lower than in China (0.68), reflecting the a distributed, less hub-centric network. Importantly, the two dominant clades sampled in Europe (ESL2.3 and ESL2.5) showed similar dissemination velocities (~600 km in 10 years; **Fig. 2e**), contrasting with clade-specific velocity differences observed in China. Together, these findings indicate that the same pathogen can exhibit fundamentally different spatial dynamics depending on underlying healthcare and patient-referral network architecture.

### Multidimensional Urban Drivers of ESL2.5 Dominance

ESL2.5’s accelerated spread in China was attributable to its disproportionate prevalence in central transmission nodes (Spearman ρ = 0.61, *p* < 0.001; **Fig. 3a**), contrasting sharply with the peripheral distribution of ESL2.4 (ρ = −0.32, *p* = 0.017; **Fig. 3b**). Consistent with this structural pattern, patient mobility data^13^ revealed that cities experiencing higher inter-city healthcare demand carried disproportionately higher burden of ESL2.5 (**Fig. 3c; See Methods**).

**Fig. 3:**
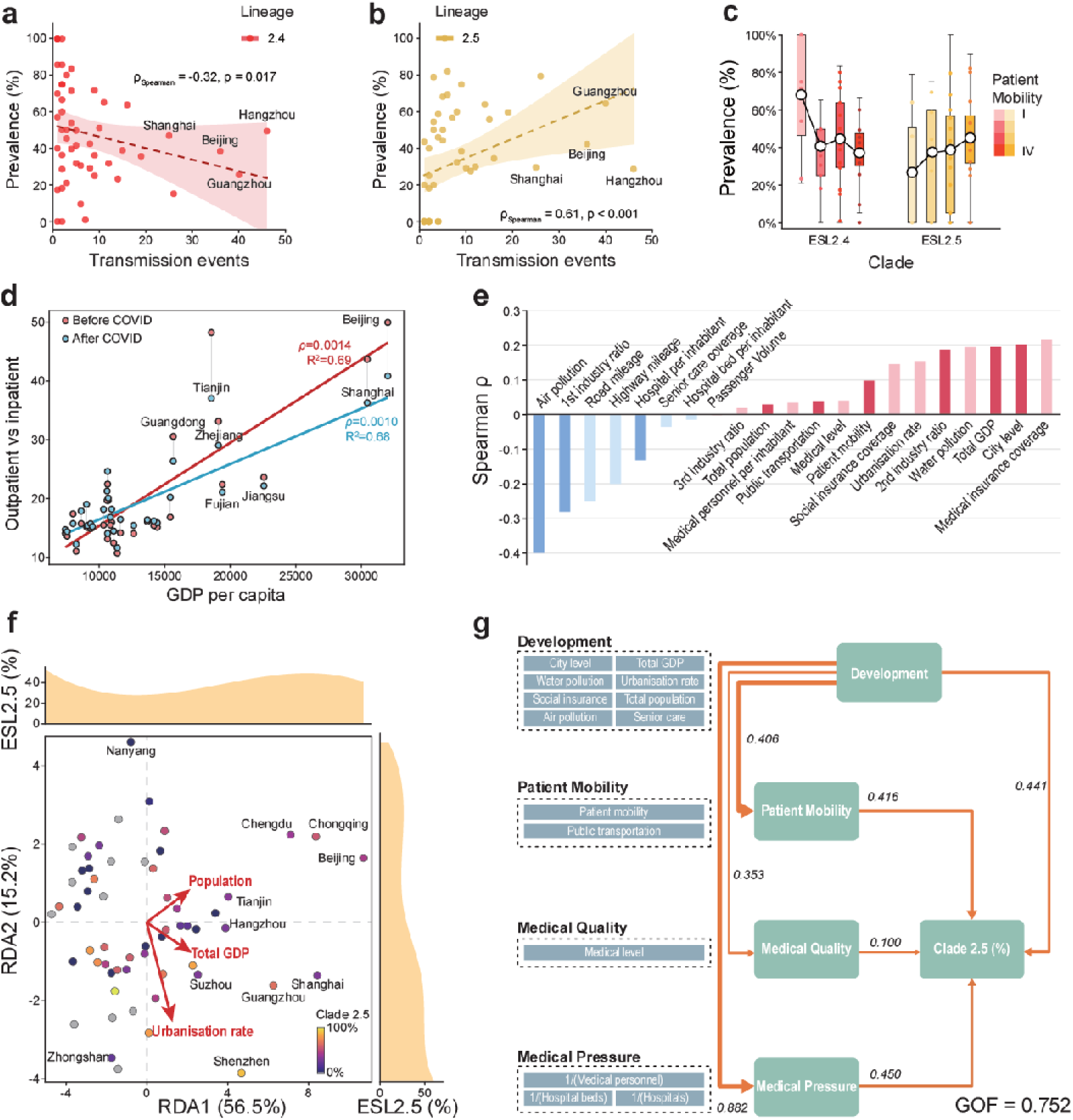
Urban determinants and healthcare-system drivers of ESL2.5 prevalence. **a-b,** Associations between city-level clade prevalence and transmission connectivity. Each point represents a city, plotted by the number of inferred transmission events and the relative prevalence of ESL2.4 (**a**, ρ = −0.32, *P* = 0.017) or ESL2.5 (**b**, ρ = 0.61, *P* < 0.001). Dashed lines show fitted Spearman correlations with shaded areas indicating 95% confidence intervals. **c,** Distribution of ESL2.4 and ESL2.5 prevalence across cities stratified by patient mobility level (I–IV). Box plots show the median and interquartile range, with individual cities overlaid as points. **d,** Association between outpatient-to-inpatient (O/I) ratio and GDP per capita across Chinese provinces before and after the COVID-19 pandemic. Each point represents a province. **e,** Spearman correlations between ESL2.5 prevalence and city-level variables. Dark-colored bars indicate variables with statistically significant correlations. **f,** Redundancy analysis (RDA) of city-level ESL2.5 prevalence constrained by urban attributes. Points represent cities, coloured by ESL2.5 relative abundance. Arrows indicate the direction and strength of explanatory variables. Marginal density plots show the distribution of ESL2.5 prevalence along each RDA axis. **g,** Structural equation model (SEM) summarizing the relationships among urban development, patient mobility, medical quality, medical pressure, and ESL2.5 prevalence. Standardized path coefficients are shown along arrows. GOF of model is indicated.

To systematically evaluate urban determinants, we integrated national healthcare and demographic datasets with city-level clade prevalence (**Table S2 and S3, See Methods**). ESL2.5 correlated positively with administrative hierarchy, gross domestic product (GDP), urbanization rate, public transit passenger volume, and medical insurance coverage. In contrast, ESL2.4 was associated with lower development indicators: air pollution levels, agricultural employment, road mileage, hospital number, and eldercare coverage (**Fig. 3e** and **Fig. S5**).

Redundancy analysis (RDA) demonstrated that aggregated city attributes explained over 70% of variance in clade prevalence (**Fig. 3f**). ESL2.5 prevalence exhibited strong positive correlations with both RDA gradients, reflecting associations with city scale (population and GDP in RDA1) and urban development indicators (urbanization and modernization in RDA2). Structural equation modeling (SEM; goodness-of-fit = 0.752) further indicated that urban development enhances ESL2.5 prevalence primarily through increased patient mobility and healthcare efficiency (**Fig. 3g**).

Notably, we observed a positive correlation between ESL2.5 prevalence and medical pressure (inverse of hospital and inpatient bed numbers per inhibitan), suggesting greater representation in cities experiencing high medical pressure, a characteristic feature of mega-cities. This interpretation was supported by elevated outpatient-to-inpatient ratios (O/I ratios) in regions with higher GDP per capita (**Fig. 3d and Table S4**), particularly in mega-cities such as Beijing and Shanghai (O/I ratio: 35–50) compared to less developed regions (O/I ratio: 10–25). These patterns suggest that ESL2.5 has undergone specialized adaptation to high-throughput urban healthcare selective pressures.

### ESL2.5 Exhibits Distinctive Gene Disruption and High-Density Mutation Patterns

To identify the genetic basis of ESL2.5’s epidemiological success, we performed pan-genome analysis, which identified a total of 36,992 pan-genes in 1,782 representative genomes (**Fig. 4a**). A key distinguishing feature was capsule conversion, shifting from predominantly KL2 in ESL2.4 to KL3 in ESL2.5.

**Fig. 4:**
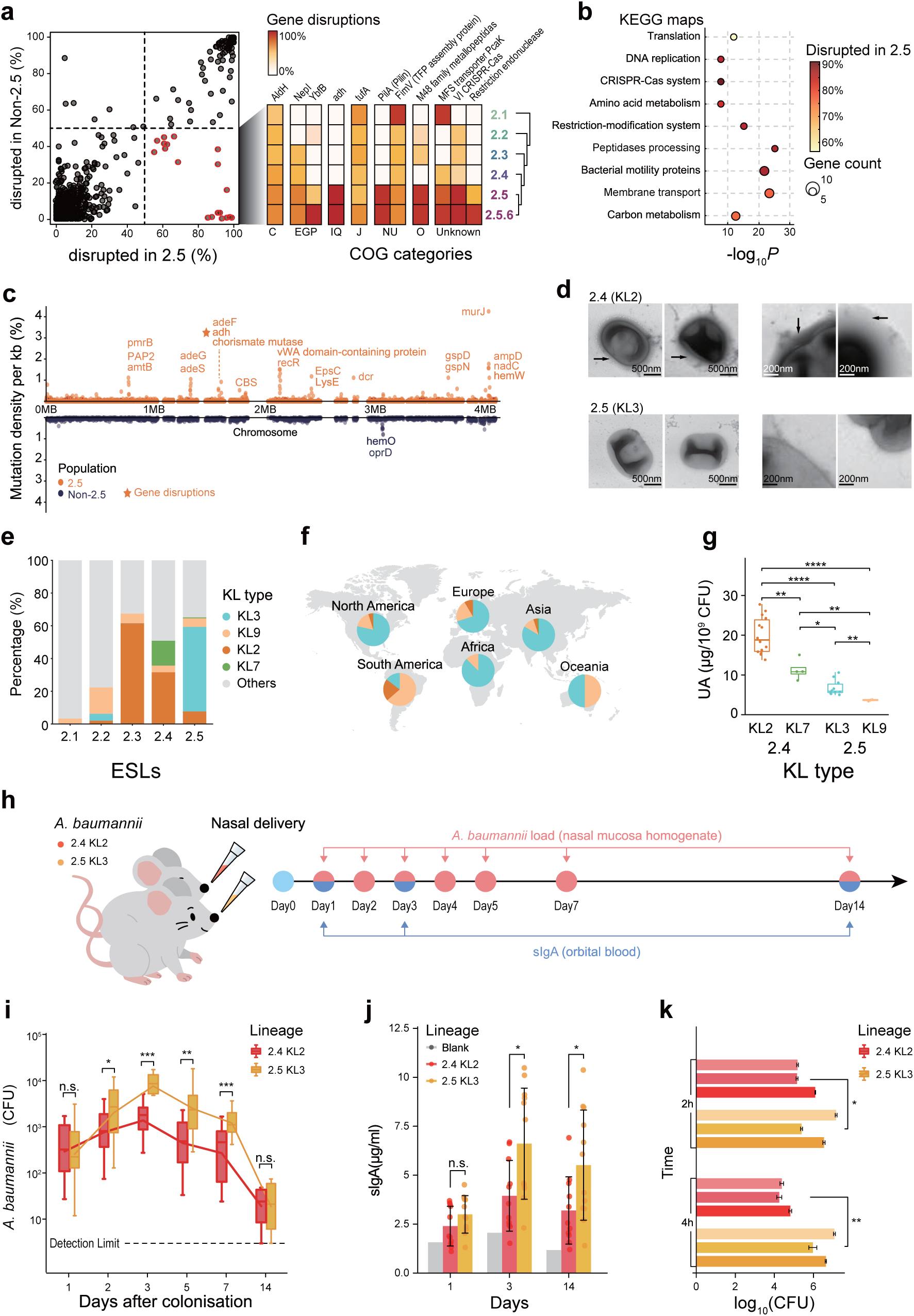
Clade-specific gene disruption reshapes host interaction. **a,** Gene-level disruption frequencies in ESL2.5 relative to non-ESL2.5 lineages. Each point represents a gene, plotted by the proportion disrupted in ESL2.5 (x-axis) versus non-ESL2.5 lineages (y-axis). Genes enriched for disruption in ESL2.5 are highlighted. The heatmap summarizes disruption frequencies across major COG functional categories for ESL sublineages. **b,** Functional enrichment of disrupted genes in ESL2.5. Bubble plot shows enriched KEGG functional categories among disrupted genes, with bubble size indicating gene count and color representing the proportion disrupted in ESL2.5. Statistical significance is shown as −log_10_*P* values. **c,** Genome-wide distribution of mutation density among non-ESL2.5 lineages and ESL2.5, with mutation hotspots highlighted by genes within. **d,** TEM images showing capsule-associated outer layer differences between ESL2.4 (KL2) and ESL2.5 (KL3) strains. Arrows indicate capsular material surrounding the bacterial cell surface. Scale bars are shown in nanometres(nm). **e,** Stacked bar plots show the relative proportions of major KL types (KL2, KL3, KL7, and KL9) across ESL2.1-2.5 lineages. **f,** Pie charts show the global distribution of KL types within non-ESL *A. baumannii*. **g,** Quantification of uronic acid (UA) content in representative KL types. Box plots show UA levels normalized to bacterial counts (μg/10^9^ CFU). Statistical significance between KL types is indicated (\**P* < 0.05; \*\**P* < 0.01; \*\*\**P* < 0.001; \*\*\*\**P* < 0.001; Mann-Whitney U test). **h,** Experimental design of the murine nasal colonization model. Mice were intranasally inoculated with ESL2.4 or ESL2.5 strains. Sampling time points for nasal bacterial load (Day1, 2, 3, 4, 5, 7, 14) and serum IgA measurements (Day1, 3, 14) are indicated. **i,** Dynamics of *A. baumannii* nasal colonization following inoculation. Bacterial loads are shown as CFU over time for ESL2.4 and ESL2.5. The dashed line indicates the detection limit. **j,** sIgA levels measured at indicated time points following nasal colonization. Box plots show sIgA concentrations for mice inoculated with ESL2.4, ESL2.5, or control groups. **k,** Adhesion of *A. baumannii* ESL2.4 and ESL2.5strains to A549 cells. Adherent bacteria were quantified at 2 h and 4 h post-infection (\**P* < 0.05; \*\**P* < 0.01; \*\*\**P* < 0.001; Mann-Whitney U test).

This structural modification is accompanied by genomic streamlining, including the disruption of 19 core genes involved in surface motility (e.g., *fimV*, *pilA*) and metabolic pathways (**Fig. 4b, Table S5**). These lesions comprised repeated frameshifts, premature stop codons, and IS-mediated truncations, validated through PCR-based sequencing (see **Methods**; **Fig. 4b** and **Fig. S6**).We also identified nine ESL2.5–specific mutational hotspots, involving efflux systems (*ade* family), lipid A/LPS modification (*pmrB*), and membrane homeostasis (*lysE*) (**Fig. 4d**), all consistent with adaptation to nutrient-poor, antimicrobial-rich hospital environments.

### ESL2.5 Demonstrates Superior Host Colonization

Transmission electron microscopy (TEM) and uronic acid content measurements confirmed that the K3 capsule was thinner than KL2 (**Fig. 4e**), potentially facilitating colonization while reducing virulence^5^. Unsurprisingly, ESL2.5 strains exhibited enhanced epithelial cell adhesion *in vitro* than ESL2.4 (**Fig. 4h**). In murine nasopharyngeal colonization models, ESL2.5 also displayed significantly higher bacterial loads over a 14-day period (**Fig. 4f**). During days 0–7, ESL2.5 burdens were approximately tenfold higher than ESL2.4 (*p* < 0.01). ESL2.5 also elicited elevated mucosal immune responses, with significantly higher sIgA concentrations on days 1 and 3 (*p* < 0.01) and sustained elevation through day 14 (**Fig. 4g**).

These findings, together with recent evidence that ESL2.5 strains exhibit lower lethality compared with ESL2.3 and ESL2.4^2^, provides a potential explanation for the apparent paradox of attenuated acute virulence coupled with superior epidemiological fitness.

### Convergent Evolution in *Klebsiella pneumoniae* ST11

The evolutionary trajectory of ESL2.5 is not an isolated phenomenon but part of a broader, taxon-independent response to the selective architecture of large-scale healthcare systems. Strikingly, we identified similar adaptive signatures in *Klebsiella pneumoniae* ST11, a phylogenetically distant nosocomial pathogen that has undergone parallel ecological specialization in China (**Table S6**). The dominant ST11-KL64 sublineage^5^, which emerged in the early 2010s, has displaced its ancestor ST11-KL47 through a pattern of rapid expansion followed by stabilization after 2017, mirroring the dynamics of ESL2.5 in *A. baumannii* (**Fig. S7a**). Phylogeographic reconstruction confirms that KL64 originated and radiated from the same mega-city hubs, Beijing and Shanghai, that serve as epicenters for ESL2.5 dissemination (**Fig. S7b**), reinforcing the central role of high-connectivity urban hospitals as crucibles of pathogen evolution.

At the genomic level, KL64 exhibits hallmark features of hospital-adapted streamlining: a serotype switch from KL47 to KL64, recurrent loss-of-function mutations in motility and surface adhesion genes (including *fimA*), and concentrated mutational hotspots in loci governing envelope integrity and stress response (**Fig. S7c-e**). KEGG pathway enrichment reveals significant convergence with ESL2.5 in the disruption of iron-acquisition systems and biofilm-associated modules, functions central to persistence in nutrient-limited, antimicrobial-rich hospital environments (**Fig. S7d**). These parallel genomic trajectories, emerging independently in two ESKAPE pathogens, provide compelling evidence that modern healthcare infrastructure imposes deterministic selective regimes that override phylogenetic history, driving convergent evolution toward colonization-optimized, transmission-efficient ecotypes.

### Simulations Validate Niche-Specific Transmission Models

The two co-circulating clades in China exhibited highly similar genome context (2043 SNP differences on average) yet contrasting ecological behaviors: ESL2.5 with enhanced upper-airway colonization and mobility-associated dissemination versus ESL2.4 with greater virulence and peripheral localization. To evaluate whether these contrasting traits recapitulate their spatial coexistence, we simulated transmission using a scale-free network representing the hierarchical Chinese healthcare system (**Fig. 5a**; **See Methods**). Each node (city) was parameterized by population size, healthcare tier, outpatient-to-inpatient (O/I) ratio, and patient mobility. Within each node, infections were modeled with distinct outpatient and inpatient contact layers. Strains were assigned two trait parameters: an inpatient colonization rate β reflecting long-term infection capacity, and an outpatient colonization rate τ associated with mobility-associated dissemination.

**Fig. 5:**
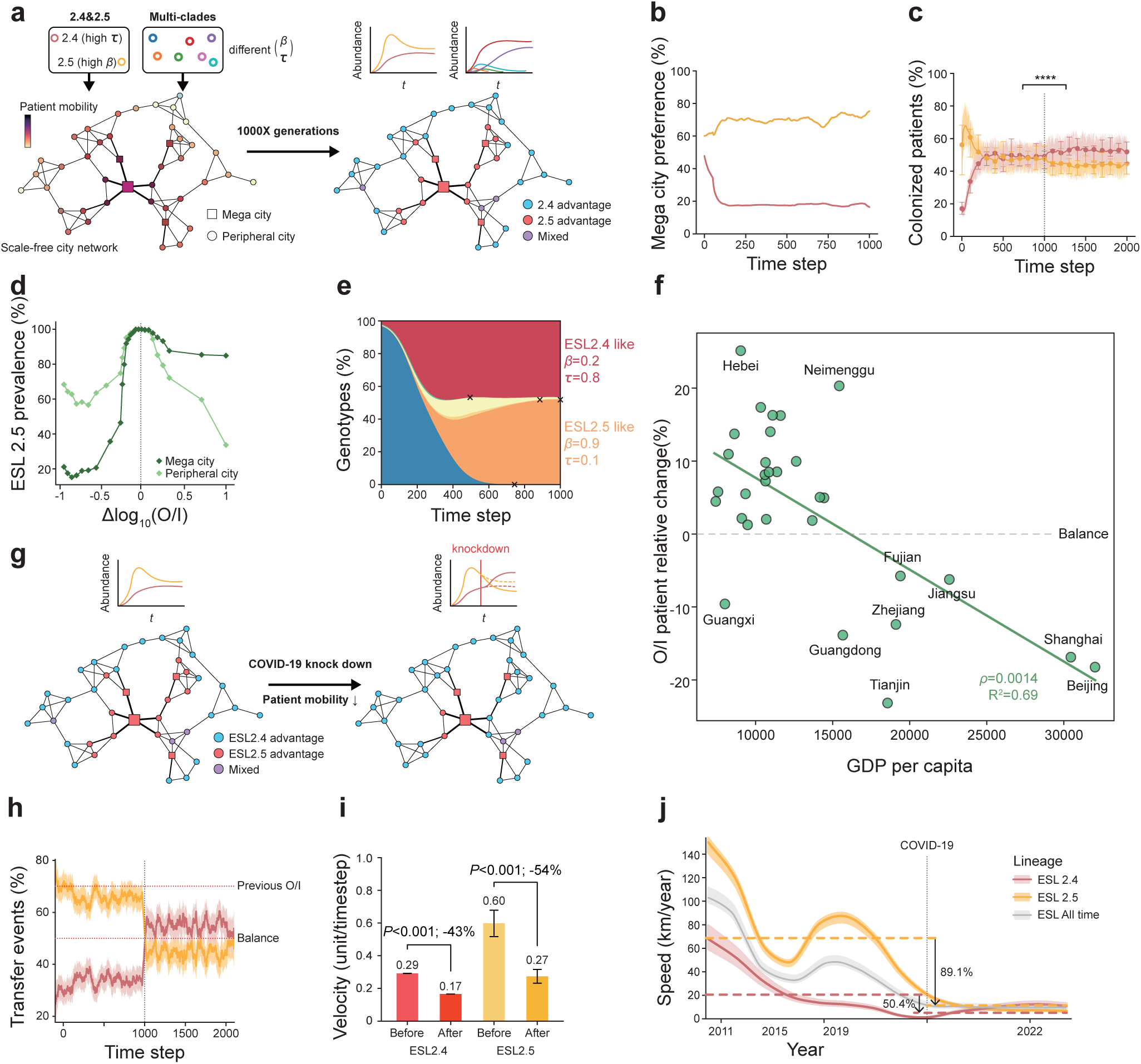
Simulation-based validation of niche-specific transmission. **a,** Schematic of the simulation framework modeling pathogen transmission on a scale-free city network representing the hierarchical Chinese healthcare system. Nodes represent cities categorized as mega-cities or peripheral cities, parameterized by outpatient-to-inpatient (O/I) ratio, and patient mobility. Transmission occurs through outpatient and inpatient contact layers. Two genotypes, representing ESL2.4 and ESL2.5, are assigned distinct inpatient (β) and outpatient (τ) colonization parameters. **b,** Temporal dynamics of genotype preference for mega-city nodes. Lines indicate the proportion of each genotype occupying mega-city nodes over simulation time. **c,** Total proportion of colonized patients over time for simulations initiated with mixed genotypes (\*\*\*\**P* < 0.001; Mann-Whitney U test). Shaded areas indicate variability across replicate simulations. The dashed line denotes the time at which COVID-19 period began. **d,** Relationship between ESL2.5 prevalence and the difference in O/I ratios between mega-city and peripheral nodes. **e,** Genotype composition over time in simulations involving multiple competing genotypes with randomly assigned (β, τ) parameter combinations. The system converges toward retention of two dominant genotypes; each associated with a distinct city class. Cross mark indicated **f,** Association between changes in provincial outpatient-to-inpatient (O/I) ratios and GDP per capita during the COVID-19 period. Each point represents a province. **g,** Simulation of pandemic-associated reductions in patient mobility. Network structure and genotype advantages are shown before and after simulated mobility suppression, mimicking non-pharmaceutical intervention effects. **h,** Predicted changes in transmission event frequencies over time under simulated pandemic conditions. Lines represent mean values across replicates. **i,** Comparison of dissemination velocities for ESL2.4 and ESL2.5 before and after simulated mobility reduction. Bars indicate mean velocities, with error bars representing variability across simulations. **j,** Time-resolved dissemination speeds of ESL2.4, ESL2.5, and all IC2 lineages inferred from empirical data. The dashed vertical line marks the onset of the COVID-19 period. Percentage reductions in dissemination speed during the pandemic interval are indicated. An interactive version of the simulated transmission network is available online at:https://naclist.github.io/naclist-portfolio/sources/ab_china_transmission_simulation/.

When two genotypes representing ESL2.4 and ESL2.5 were introduced, the system consistently converged to stable spatial segregation. ESL2.5 dominated highly connected mega-city nodes, while ESL2.4 dominated low-connectivity peripheral nodes, regardless of starting frequencies (**Fig. 5b, c**). This pattern recapitulated the observed geographic distributions (**Fig. 3a, b**) and resulted from higher τ values in ESL2.5, which conferred significantly faster dissemination rates through outpatient networks.

Random perturbation of model parameters identified the O/I ratio as the most influential node-level determinant of coexistence. Long-term co-circulation emerged only when mega-cities and peripheral cities differed in O/I ratios; equalizing this parameter collapsed the system to single-genotype dominance (**Fig. 5d**). To further test niche partitioning, we simulated competition among 10–100 genotypes with randomly assigned (β, τ) values. When run to equilibrium, the network consistently retained only two genotypes, each specialized to one node class, a robust “one niche, one genotype” outcome (**Fig. 5e**). These results demonstrate that healthcare infrastructure alone creates distinct ecological niches capable of maintaining phenotypically divergent lineages.

### COVID-19 Pandemic Illuminates Transmission Dependencies and Systemic Vulnerabilities

The model made a testable prediction: if ESL2.5’s success depends on high outpatient mobility while ESL2.4 relies on inpatient networks, then disruptions differentially affecting these pathways should produce clade-specific epidemiological responses. The COVID-19 pandemic provided precisely such a natural experiment. Provincial healthcare statistics showed pronounced O/I ratio reductions in high-GDP regions (**Fig. 5f**), driven by large declines in outpatient visits due to non-pharmaceutical interventions (NPIs), whereas inpatient transfers were less affected.

We incorporated these empirically observed shifts into our simulation model (**Fig. 5g**). The modified model predicted reduced prevalence and dissemination speed for ESL2.5, which relies heavily on outpatient-mediated mobility (**Fig. 5c, h**). In contrast, ESL2.4, linked to inpatient networks, was predicted to experience only a moderate decline in velocity and transiently increased in relative prevalence during the NPI period (**Fig. 5i**).

Real-world surveillance closely matched these predictions (**Fig. 5j**). Between 2020 and 2023, dissemination speed attributable to ESL2.5 dropped by 89.1% (65.2 to 7.1), compared with a smaller 50.4% reduction for ESL2.4 (20.4 to 10.3). The relative frequency of ESL2.5 also decreased to 45% during the NPI period, before rebounding to 73% in 2023 (**Fig. 1b**). The pandemic perturbation thus served as an unintended experimental manipulation, confirming that lineage fitness is tightly constrained by healthcare-network structure, and that phenotypic specialization to specific transmission pathways determines epidemiological success.

## Discussion

This study demonstrates that healthcare infrastructure operates as an underappreciated yet powerful evolutionary force shaping the dissemination and adaptation of multidrug-resistant *A. baumannii*. By integrating large-scale genomics with detailed healthcare and mobility data, we demonstrate that structural features of healthcare systems, particularly network connectivity and patient flow patterns, impose ecological constraints that systematically shape pathogen population structure, niche preference, and transmission potential. This reframes pathogen evolution beyond a singular focus on antibiotic pressure to include infrastructural selection as a central, actionable evolutionary driver.

A central insight of this work is that clade replacement in Chinese *A. baumannii* is driven by transmission efficiency rather than resistance superiority. Despite near-equivalent resistomes between ESL2.4 and ESL2.5, ESL2.5 expanded more rapidly and broadly, both temporally and spatially. This finding exemplifies how long-standing endemic lineages can be displaced by more mobile, transmission-optimized variants when healthcare connectivity intensifies. ESL2.4 represents a deeply rooted lineage with decades of persistence, consistent with adaptation to lower-connectivity inpatient environments where prolonged infection and virulence may be selectively advantageous. In contrast, ESL2.5, emerged only in the late 1990s, exhibits hallmarks of a “transmission-optimized” lineage, with a thinner K3 capsule, enhanced epithelial adhesion, and superior upper respiratory tract colonization. These all favor silent, high-throughput dissemination across outpatient-dense networks.

Critically, this dichotomy is not unique to *A. baumannii*. We observe strikingly parallel genomic and epidemiological trajectories in *Klebsiella pneumoniae* ST11-KL64, which has similarly displaced its KL47 ancestor through serotype switching, loss-of-function mutations in fimbrial genes, and expansion from mega-city hubs, further evidence that modern hospital ecosystems select predictably for colonization-efficient variants. Together, these findings recast the classical virulence-transmissibility trade-off^14^ not as a simple inverse correlation, but as a spatially explicit ecological partitioning across heterogeneous healthcare landscapes^15,16^.

Our reconstruction of transmission networks reveals that Chinese healthcare operates as a scale-free, hub-dominated system in which a small number of mega-city hospitals disproportionately shape national pathogen dynamics^5^. Such architectures are common in developing regions, driven by inadequate primary care infrastructure^17,18^, and known to amplify selection for traits that enhance dissemination through hubs while penalizing strategies optimized for localized persistence^19^. ESL2.5’s preferential occupancy of these hubs, coupled with its higher migration velocity along identical network routes, indicates that it possesses phenotypic features enabling more efficient exploitation of outpatient-mediated transmission. Importantly, when the same pathogen was examined in Europe, where referral systems are flatter and inter-regional flows less hub-centric, clade-specific velocity differences disappeared. This contrast provides compelling evidence that healthcare architecture, rather than intrinsic lineage properties alone, determines which evolutionary strategies are favored.

The strong association between ESL2.5 prevalence and indicators of urbanization, mobility, and medical pressure suggests that high-throughput healthcare environments impose distinctive selective pressures. Elevated outpatient-to-inpatient ratios, characteristic of Chinese mega-cities, likely favor lineages capable of persistent colonization with limited overt disease, enabling silent carriage across large patient populations. Similar associations between healthcare intensity and pathogen evolution have been proposed^20^, where shortened lengths of stay and high patient turnover select for colonization efficiency rather than acute virulence. Our data provides genome-resolved evidence for this process in *A. baumannii*.

At the genomic level, ESL2.5 exhibits a striking pattern of capsule conversion, gene disruption, and mutational clustering affecting motility, membrane transport, immune defense systems, and metabolic flexibility. Rather than representing random decay, these lesions appear selectively structured, arising early during clade divergence and repeatedly targeting similar functional modules. Genome streamlining via loss-of-function mutations has been increasingly recognized as an adaptive strategy in stable, resource-limited, or host-associated environments^21^. The convergence of similar disruption patterns repeatedly documented from chronic infections^22^ and long-term hospital colonization^23^ strengthens the argument that these are generalizable responses to hospital selection rather than lineage-specific anomalies.

Functionally, the phenotypic consequences of this genomic remodeling are profound. Reduced type IV pilus expression and motility in ESL2.5 are accompanied by enhanced epithelial adhesion and superior nasopharyngeal colonization, both *in vitro* and *in vivo*. This trade-off aligns with theoretical expectations that loss of motility can increase surface attachment and persistence, particularly in mucosal niches^24^. The observed attenuation of acute virulence, together with heightened colonization capacity, explains how highly successful lineages can spread widely without causing proportionally higher mortality.

The integration of empirical data with mechanistic simulations represents a key advance of this work. By explicitly modeling outpatient and inpatient transmission layers within a scale-free healthcare network, we show that stable coexistence of divergent lineages emerges naturally from infrastructure-imposed niche partitioning. The robustness of the “one niche, one genotype” outcome across wide parameter ranges argues against historical contingency or founder effects as primary explanations. This finding resonates with patterns observed in microbial communities where resource specialization enables coexistence of otherwise competing taxa^25^, supporting a deterministic framework in which healthcare systems generate reproducible evolutionary outcomes.

The COVID-19 pandemic provided an unplanned yet powerful perturbation that validated our model’s core predictions. The disproportionate collapse of ESL2.5 dissemination following reductions in outpatient mobility offers rare causal evidence linking lineage fitness to specific healthcare pathways. Comparable pandemic-associated shifts have been reported for respiratory viruses and some bacterial pathogens, but rarely with this level of genomic and mechanistic resolution^26^. The rapid rebound of ESL2.5 following relaxation of NPIs further underscores how external perturbations to healthcare mobility can fundamentally alter competitive dynamics between co-circulating ecotypes.

While our study demonstrates stable coexistence of ESL2.4 and ESL2.5 through niche specialization, this ecological partitioning also creates conditions for potential evolutionary convergence that could pose substantial clinical threats. *A. baumannii* exhibits natural competence and documented recombination rates between co-circulating lineages. High prevalence and geographic overlap of these clades in tertiary hospitals, particularly in mega-cities where both lineages co-circulate, provides opportunities for horizontal gene transfer, recombination, and competitive selection that could yield hybrid strains combining ESL2.4’s virulence advantages with ESL2.5’s transmission efficiency, mirroring the convergence of virulence-resistance in *K. pneumoniae*^5^.

Several limitations warrant consideration. While our sampling is large and spatially broad, it remains uneven in time and geography; local hospital practices and undocumented patient flows could generate unmeasured heterogeneity. Second, functional assays were performed on representative isolates; although convergence across genomes and taxa supports generality, broader phenotype panels including environmental persistence, competition assays, and patient-level carriage studies will strengthen causal inference. Third, the trade-off interpretation relies on inference from correlated genomic and phenotypic signatures. Future studies integrating patient-level mobility data, electronic health records, and antimicrobial usage profiles will be critical for refining the predictive accuracy of infrastructure-based models.

In conclusion, our findings redefine how pathogen evolution should be conceptualized in modern healthcare systems. Rather than selecting uniformly for resistance and virulence, hospital networks create heterogeneous ecological niches that favor distinct, predictable evolutionary strategies. Recognizing healthcare infrastructure as a driver of pathogen diversification has immediate translational implications. Surveillance programs should stratify risk by hospital connectivity and patient flow, not solely by resistance profiles. Interventions that modulate outpatient throughput or referral patterns may have evolutionary consequences as significant as antimicrobial stewardship. More broadly, as healthcare systems continue to expand and intensify globally, understanding and potentially redesigning their ecological footprint may become a critical component of controlling the next generation of hospital-adapted pathogens.

## Methods

### Bacterial culture and whole-genome sequencing

We collected 1042 clinical isolates of *Acinetobacter baumannii* from 20 tertiary hospitals distributed across 15 provinces in China between 2018 and 2024 (**Table S1**). All isolates were resuscitated from −80°C storage and subcultured under standardized laboratory conditions at the Second Affiliated Hospital of Soochow University. Species identity was confirmed using matrix-assisted laser desorption/ionization time-of-flight mass spectrometry (MALDI-TOF MS; bioMérieux, France) with a log score threshold of ≥2.0 for species-level identification.

Genomic DNA was extracted using the QIAamp DNA Mini Kit (Qiagen, Hilden, Germany) following the manufacturer’s protocol. Sequencing libraries were prepared using the Nextera XT DNA Library Preparation Kit (Illumina, San Diego, CA) and subjected to paired-end 250 bp whole-genome sequencing on an Illumina NovaSeq 6000 platform.

### Genomic analysis

Raw sequencing reads were quality-filtered using BBduk2 in BBmap package with parameters “ordered=t ref=adapters ktrim=r overwrite=t refstats=PE.refstats qout=33 k=25 mink=13 minlength=23 tbo=t entropy=0.75 entropywindow=25 mininsert=23 maxns=2 trimq=6 qtrim=rl”. Draft genomes were assembled and quality-controlled using EToKi assemble v.1.3^27^.

To provide global context, we retrieved 3,981 assembled genomes of A. baumannii from the NCBI RefSeq database (accessed March 2024) with assembly quality filters requiring genome completeness >95% and contamination <5% as assessed by FetchMG (http://motu-tool.org/fetchMG.html). The final dataset comprised 5,023 genomes collected between 1999 and 2024 from 73 cities across 27 provinces in China. For comparative analysis with European strains, we downloaded 2,800 publicly available *A. baumannii* genomes from 12 European countries sampled between 2010 and 2023. For comparative analysis with the United States, we analyzed 5,681 publicly available genomes sampled between 2005 and 2024. Lineage assignments for genomic samples were determined using Capybara^2^. For interspecies comparative analysis, we downloaded 847 ST11 *Klebsiella pneumoniae* genomes, including both KL47 and KL64 capsular types, as previously described by Hu et al.^5^.

### Urban metadata

To investigate the relationship between *A. baumannii* dissemination patterns and healthcare infrastructure, we integrated comprehensive city-level socioeconomic, demographic, and healthcare indicators for all sampled cities. Data on healthcare accessibility and intercity patient mobility were obtained from the hierarchical classification of healthcare supply-demand networks and urban medical mobility patterns constructed by Zhao et al. (2025) ^13^ based on national Baidu migration data representing over 200 million individual movement events. Additional urban attributes were extracted from the China Statistical Yearbook 2023 (National Bureau of Statistics, https://www.stats.gov.cn) ^28^ and the China Health Statistical Yearbook 2013–2023 (National Health Commission, https://www.nhc.gov.cn/mohwsbwstjxxzx/tjnj) ^29^. Variables included: administrative hierarchy (provincial capital, prefecture-level city, or county-level city), gross domestic product (GDP) and GDP per capita, total population and urbanization rate (proportion of urban residents), public transport passenger volume (annual bus and metro ridership), numbers of medical insurance and social security participants (as a proxy for healthcare access), air quality index (annual PM2.5 concentrations), total road mileage, drainage pipeline coverage (as infrastructure proxy), numbers of hospitals and inpatient beds, numbers of healthcare workers (physicians and nurses per 1,000 population), eldercare coverage (nursing home beds per 1,000 elderly population), outpatient visits and inpatient admissions (used to calculate outpatient-to-inpatient [O/I] ratios), and average hospital length of stay. Medical pressure was defined as the inverse of hospital beds per capita and inpatient beds per capita, representing the ratio of healthcare demand to supply.

### Construction of time-resolved phylogeny

We reconstructed a time-calibrated phylogeny using recombination-free SNPs from conserved genomic regions following established procedures. All genomes were aligned to the reference strain MDR-TJ (GCF_000187205.2; isolated from a Chinese hospital in 2013) using EToKi align. Recombinant regions were detected and masked using RecHMM v.08d7382^30^ and ClonalFrameML v.1.13^31^, which yielded almost identical results. A maximum-likelihood (ML) phylogeny was then inferred with IQ-TREE2 v.2.2.2.6 ^32^ under a GTR+G substitution model. The temporal signal was evaluated by regression of root-to-tip distances against sampling dates using TempEst v.1.5.3^33^, yielding a correlation coefficient (R) of 0.13 (P < 0.001).

Time-scaled phylogenies were inferred using TreeTime v.0.11.3^34^ with a relaxed molecular clock model (coefficient of variation = 0.3), a coalescent skyline tree prior, and confidence intervals estimated from 100 bootstrap replicates of the input alignment. To validate temporal inference, we performed independent dating analysis using BEAST2 v.2.6.6^35^ under both strict and relaxed (uncorrelated lognormal) clock models with a Bayesian skyline coalescent prior, sampling every 10,000 steps for 50 million generations after 10% burn-in. Effective sample sizes (ESS) for all parameters exceeded 200. Divergence time estimates and evolutionary rate estimates from TreeTime and BEAST2 agreed within 95% highest posterior density intervals. Given computational efficiency and robustness to model violations in large datasets, TreeTime was used for the full 5,023-genome dataset. Ancestral state reconstruction for geographic locations was performed using TreeTime’s mugration module with a constant transition rate model and discrete trait optimization.

### Transmission inference

Transmission dynamics were inferred using the trait reconstruction module of TreeTime, which estimates the posterior probability of the time to the most recent common ancestor (tMRCA) for each node. A transmission event was defined as any branch connecting two nodes assigned to different geographic locations. For each sequence pair, the distribution of divergence times was used to infer potential transmission pathways. Each branch thus represented a putative transmission link, providing an opportunity for cross-regional migration. For a branch *e* = (*u*–*v*), where the parent (*u*) and descendant (*v*) nodes were inferred to differ in geographic state, the conservative transmission velocity (*V_e_*) was defined as:

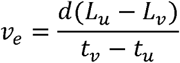

where *t_(u)_* and *t_(v)_* denote the inferred node ages estimated from the dated phylogeny, *L_(u)_* and *L_(v)_* represent the most probable geographic locations of nodes *u* and *v*, and *d*(*L_(u)_*- *L_(v)_*) is the geographic distance between these locations. To assess robustness, we constructed two transmission matrices: one weighted by the ancestral location probabilities of each node, and another restricted to nodes with high-confidence (>70%) ancestral state assignments. Cities were treated as nodes in a directed transmission network, and the community structure of this network was detected using the Louvain modularity algorithm implemented in Gephi v.0.10.1^36^. Multiple resolution parameters were tested to explore potential multilevel community partitions. Using the default resolution of 1.0, the directed network was partitioned into six major modules, representing nationwide transmission communities, with an overall modularity (Q) value of 0.59. Power-law scaling was used to evaluate scale-free network metrics both within individual modules and across the overall network.

To quantify geographic constraints on dissemination, we classified each inferred transmission event as within-module (*w*) or between-module. Geographic distances between city pairs were grouped into 100-km bins, and the proportion of within-module events in each bin (ratio of *w*) was computed to normalize for uneven event counts across distance classes. To reduce stochastic fluctuations, we applied LOESS smoothing and generated 95% confidence intervals via 1,000 bootstrap replicates.

The transmission boundary was defined as the smallest distance at which the smoothed ratio of w declined to 0.5, representing the scale at which within-module and between-module events occurred with equal probability. Across all bootstrap iterations, this threshold consistently appeared near ~800 km, indicating a transition point beyond which geographic barriers substantially weakened.

### Multidimensional analysis of city attributions

To systematically evaluate the influence of socioeconomic and healthcare-related factors on the prevalence of ESL2.5, we compiled 22 city-level variables from the China Statistical Yearbook (2023)^28^ and the China Health Statistical Yearbook (2013–2023)^29^ and assessed their associations with city-level ESL2.5 relative abundance using Spearman rank correlations.

For each variable, the Spearman ρ coefficient was calculated and ranked by sign and significance to distinguish suppressing factors (negative correlations) from promoting factors (positive correlations). Variables exhibiting significant correlations were retained as candidate predictors for subsequent RDA and SEM analyses (**Table S2**).

To identify key drivers of spatial differences in ESL2.5 abundance, redundancy analysis (RDA) was performed using the vegan R package^37^, with city-level ESL2.5 relative abundance as the response variable and the significantly associated socioeconomic and healthcare variables as explanatory matrices. All explanatory variables were centered and standardized prior to analysis. The significance of the RDA model was evaluated using 9,999 permutation tests, and the first two constrained axes (RDA1 and RDA2) were extracted to visualize variable gradients and city positioning. Arrow orientation indicated the direction of increasing variable gradients, and arrow length reflected explanatory strength. Population size, GDP, and urbanization rate emerged as the predominant contributors to the spatial heterogeneity of ESL2.5.

To further quantify the multilayered effects of urban development, healthcare resources, healthcare quality, and population mobility on ESL2.5 prevalence, we constructed an extended structural equation model (SEM). Based on functional groupings of urban attributes, variables were organized into four latent components: Development, Medical capacity, Medical quality, and Patient mobility. Each component was represented by a standardized composite score derived from its representative indicators. The SEM specified a series of linked linear equations to construct causal pathways, estimating the following relationships:

1. Medical capacity as a function of Development;
2. Medical quality as a function of Development and Medical capacity;
3. Patient mobility as a function of Development;
4. ESL2.5 relative abundance as a function of Development, Medical capacity, Medical quality, and Patient mobility.

All variables were z-score standardized before modeling. Each path was fitted using ordinary least squares (OLS), and standardized coefficients were extracted. Model fit was summarized using the geometric mean of R² values across component equations. To assess robustness, P values were reported for each pathway, enabling evaluation of multilevel mechanisms through which urban attributes influence ESL2.5 prevalence via healthcare systems or mobility-related processes. The SEM revealed that Development exerted a strong positive effect on ESL2.5 dissemination by increasing Patient mobility, whereas Medical capacity showed a significant suppressive effect. The final model achieved a GOF of 0.752, indicating high explanatory power and stability.

### Pangenome construction and mutation density estimation

All genomes were annotated using Prokka v.1.14.6^38^ with default parameters and the --kingdom Bacteria flag. Orthologous gene families were assigned using eggNOG-mapper v.1.0.5^39^ based on the eggNOG 5.0 database, with DIAMOND v.2.0.15 used for homology searches (E-value threshold 10^-5^). The complete pangenome was reconstructed using PEPPAN v.1.0.5^40^ with parameters, which performs iterative clustering of coding sequences and identifies potential pseudogenes.

Putative gene disruptions (pseudogenes) were classified into three categories based on structural defects: (1) Truncated genes: fragments <120 bp or lacking canonical start (ATG, GTG, TTG) or stop (TAA, TAG, TGA) codons; (2) Frameshift mutations: coding sequences with lengths not divisible by three, causing translational reading-frame shifts; (3) Nonsense mutations: premature stop codons within the first 90% of the expected coding sequence, verified by alignment to intact homologs. A total of 1249 pseudogenes were identified across the pangenome. To assess the contribution of insertion sequences (IS elements) to gene disruption, we identified IS elements using ISFinder^41^ (https://isfinder.biotoul.fr/) and defined IS-mediated disruptions as pseudogenes with IS elements located within the coding regions. PCR amplification and Sanger sequencing were performed to validate a subset of 15 predicted disruptions distributed across different functional categories, using primers designed to span the disrupted regions with 200-300 bp flanking sequence and sequenced bidirectionally using the ABI 3730xl platform.

To trace the evolutionary history of gene disruptions, ancestral gene states (intact, disrupted, or absent) were reconstructed across the time-calibrated phylogeny using TreeTime’s ancestral sequence reconstruction module. For each pseudogene, we identified the phylogenetic node at which the disruption most likely arose based on maximum posterior probability. Disruptions were classified as: (1) early clade-defining events if they occurred at or near the root of ESL2.5; (2) parallel losses if the same gene was independently disrupted in multiple sublineages; (3) recent losses if they occurred in terminal branches or shallow clades. The temporal distribution of disruption events was visualized by mapping disruption times onto the dated phylogeny.

To identify high-mutation-density regions (mutational hotspots), we aligned all genomes to the MDR-TJ reference using EToKi align module, which internally calls minimap2 v.2.26^42^ with preset parameters -ax asm5. SNP density was calculated in 1-kb sliding windows with 500-bp step size across the genome. Mutational hotspots were defined as windows with SNP density exceeding the 95th percentile. Each SNP was annotated for genic context (intergenic, synonymous, nonsynonymous, nonsense).

To identify functional pathways enriched among fragmented genes, all differentially disrupted coding sequences were converted to KEGG Orthology^43^ (KO) identifiers. Using species-specific KEGG annotations, gene–pathway mappings were constructed, and enrichment significance was evaluated using adjusted *p* values (<0.05). For each enriched pathway, enrichment factors were computed and visualized according to pathway categories to characterize functional biases in pseudogene accumulation.

For comparative analysis with *K. pneumoniae* ST11, we performed analogous pangenome construction, pseudogene identification, and mutational hotspot detection using the 847 ST11 genomes, following identical protocols. Functional convergence between *A. baumannii* ESL2.5 and *K. pneumoniae* ST11-KL64 was assessed by comparing enriched KEGG pathways and identifying shared disrupted gene families.

### Outer layer component identification

Single colonies were inoculated into LB broth under sterile conditions and incubated overnight at 37 °C with shaking at 180 rpm. The cultures were then diluted 1:100 into fresh LB medium and grown at 37 °C to mid-exponential phase. Bacterial cells were harvested by low-speed centrifugation and washed three times with phosphate-buffered saline (PBS). The cell pellets were immediately resuspended in 2.5% (v/v) glutaraldehyde and gently mixed to ensure uniform fixation, followed by storage at 4 °C.

Fixed samples were washed three times with PBS and post-fixed with 1% (w/v) osmium tetroxide at room temperature for 1 h under light-protected conditions. Samples were subsequently washed twice with PBS and twice with deionized water to remove residual phosphate. En bloc staining was performed using 2% (w/v) aqueous uranyl acetate for 1 h at room temperature.

After staining, samples were dehydrated through a graded acetone series (50%, 70%, 80%, 90%, and 100%) and gradually infiltrated with resin embedding mixtures prior to polymerization. Resin blocks were trimmed and sectioned into ultrathin sections, which were examined using a transmission electron microscope. Imaging was performed after vacuum stabilization, voltage ramping, and beam alignment, and images were acquired in CCD mode.

Following the identification of pronounced differences in the outer layer under transmission electron microscopy, capsular polysaccharides were extracted as previously described^5^. Briefly, *A. baumannii* strain cultures corresponding to approximately 10^9^ colony-forming units (CFU) were harvested and resuspended in 1% (w/v) Zwittergent 3-14 detergent (Sangon Biotech, China, Cat. No. A610552) in 100 mM citric acid, after which the polysaccharides were precipitated using ethanol.

Capsular polysaccharide levels were quantified using a modified phenol–sulfuric acid assay, as previously described^44^. In brief, the precipitated polysaccharide samples were resuspended in 200 μL of ddH_2_O and mixed with 400 μL of 5% (w/v) phenol, followed by the addition of 2 mL of concentrated sulfuric acid. After incubation at room temperature for 30 minutes, absorbance was measured at 490 nm. Polysaccharide concentrations were determined from a glucose standard curve.

Additionally, the capsular polysaccharide was quantified using a uronic acid assay as previously described^5^. In brief, precipitated polysaccharide samples were incubated with 0.0125 M sodium tetraborate in sulfuric acid, then reacted with 0.125% (w/v) carbazole in absolute ethanol. Absorbance was measured at 530 nm, and the uronic acid content was quantified using a D-glucuronic acid standard curve.

### Mouse nasal colonization assay

Frozen *A. baumannii* isolates stored at −80 °C were streaked on Luria-Bertani (LB) agar plates and incubated at 37 °C for recovery. A single colony was inoculated into 5 mL LB broth and cultured overnight at 37 °C with shaking at 180 rpm. The overnight culture was diluted 1:1000 into 40 mL fresh LB and incubated until mid-logarithmic phase (OD_600_ = 0.5-0.8). Cells were washed twice and resuspended in phosphate-buffered saline (PBS) to a final concentration of OD_600_ = 0.3±0.05. For nasal colonization, 40 µL of the bacterial suspension was slowly administered intranasally (20 µL per nostril) to anesthetized mice to simulate upper respiratory tract colonization. Control mice received 40 µL PBS. The initial bacterial inoculum was quantified by serial dilution and plating. Samples were collected at 1, 2, 3-, 5-, 7-, and 14-days post-inoculation. Blood was obtained by retro-orbital bleeding, and nasal mucosa tissues were excised and transferred into 1 mL PBS containing 1 mm glass beads. For colony enumeration, tubes containing nasal tissues were processed using a SCIENTZ-48 tissue grinder at 65 Hz, followed by immediate cooling on ice. 100 µL of homogenate was added to 900 µL PBS and incubated for 12 h at 37 °C, after which serial dilutions were plated for bacterial counting.

### Measurement of secretory immunoglobulin A and adhesion

The concentration of secretory immunoglobulin A (sIgA) in mouse serum was quantified using an ELK1549 ELISA kit (Kelu Biotechnology, Wuhan, China). Mice were sacrificed at 1-, 3-, and 14-days post-infection, and blood samples were collected via retro-orbital puncture. All reagents were equilibrated to room temperature prior to use. Standards were serially diluted to generate a calibration curve. Serum samples were diluted 1:10 and added to antibody-coated microplates, followed by incubation at 37 °C. Biotin-conjugated detection antibodies and HRP-labeled secondary reagents were sequentially added and incubated for 50 min each at 37 °C, with washing between steps. The colourimetric reaction was developed with TMB substrate at 37 °C in the dark, terminated with stop solution, and the absorbance was measured at 450 nm. sIgA concentrations were determined according to the standard curve.

Bacterial adhesion to epithelial cells was assessed using the human lung carcinoma cell line A549. Cells were maintained in Dulbecco’s modified Eagle’s medium (DMEM) supplemented with 10% fetal bovine serum and cultured at 37 °C with 5% CO_2_. Prior to infection, A549 cells were seeded into tissue culture plates and grown to confluence.

Overnight bacterial cultures of *A. baumannii* ESL2.4 and ESL2.5 strains were washed and resuspended in cell culture medium. After incubation for 2 h or 4 h, non-adherent bacteria were removed by washing the monolayers with phosphate-buffered saline. Host cells were subsequently lysed, and adherent bacteria were enumerated by plating serial dilutions on agar plates. All assays were performed with 3 independent replicates.

### Simulation model of intercity competition

To evaluate how heterogeneity in healthcare hierarchies’ shapes selection and migration, we developed a multi-genotype Wright–Fisher simulation across 100 cities interconnected by asymmetric mobility. Eight genotypes were defined by complementary parameters τ (outpatient colonization ability) and β (inpatient infection ability, β = 1 − τ), representing alternative dissemination strategies. Thirty-five cities were designated as metropolitan hospitals and 65 as peripheral hospitals. Each city maintained an effective population size of *N* = 1000, evolving for 1000 generations.

Genotype fitness depended on the match between τ/β values and city type, with selection strength *s* = 0.1. Each generation comprised (1) within-city Wright–Fisher reproduction with drift and selection, followed by (2) intercity migration shaped by hierarchical referral preference (peripheral-metro bias) and spatial decay modeled as:

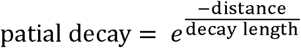

Migrants replaced random individuals in the destination population. Initial infection was seeded in 15% of cities at 5% prevalence, with infected individuals evenly split between high-τ and high-β genotypes. Two analytical modes were supported: a basic competition scenario and a public-health perturbation scenario mimicking COVID-like interventions with baseline, restriction, and recovery phases. Outputs included weighted *F_ST_*, genotype frequency variance, metro–peripheral divergence and proportion of specialist genotypes (τ > 0.6 or τ < 0.4). For genotype transferred between cities i and j, the effective dissemination distance (*d*) is defined as:

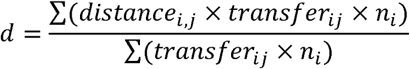

where distance_i,j_ denotes the geographic distance between cities i and j, transfer_i,j_ represents the number of inferred transmission events, and n_i_ represents the sample size of the source city.

Sensitivity analyses varied mobility ratio μ, selection strength *s*, spatial decay length, and numbers of metropolitan cities, with 10 replicates per parameter set.

### Statistical analysis

All statistical analyses were performed in R v.4.3.1 or Python v.3.10 unless otherwise specified. For comparisons between two groups, parametric (Student’s *t*-test) or non-parametric (Mann-Whitney U test) tests were applied. For multiple group comparisons, one-way ANOVA or Kruskal-Wallis tests were used, followed by appropriate post-hoc tests. Time-series comparisons in animal experiments employed two-way ANOVA with factors for strain and timepoint, followed by Tukey’s post-hoc tests. Categorical associations were evaluated using Fisher’s exact test or Chi-square test. Correlation analyses used Pearson correlation for normally distributed continuous variables or Spearman rank correlation for non-normal or ordinal data. For multiple hypothesis testing, false discovery rate (FDR) was controlled using the Benjamini-Hochberg procedure, with adjusted P-values (q-values) reported where appropriate.

## Lead contact

Further information and requests for resources and reagents should be directed to, and will be fulfilled by, the lead contact, Zhemin Zhou.

## Data and code availability

All sequencing data, figures, and analysis scripts are publicly available at GitHub (https://github.com/Zhou-lab-SUDA/Genomic_China_Acinetobacter). Assembled genome sequences have been deposited in the China National Center for Bioinformation (CNCB) under accession numbers PRJCA055264, also available in the GitHub repository and FigShare. Associated metadata are accessible via the GitHub repository, and the PathoBase database (https://pathobase.iotabiome.com/) under the corresponding project identifiers.

All scripts used for genomic analyses and figure generation are available at GitHub (https://github.com/Zhou-lab-SUDA/Genomic_China_Acinetobacter). https://naclist.github.io/naclist-portfolio/sources/ab_china_dynamics/

Any additional information required to reanalyze the data reported in this paper is available from the lead contact upon request.

## Acknowledgments

The project was primarily supported by the National Natural Science Foundation of China (82530102, 32170003, 32370099). This work was also supported by the Provincial-level Talent Program for National Center of Technology Innovation for Biopharmaceuticals (NCTIB2024JS0101), the Natural Science Foundation of Jiangsu Province (BK20243008), the Suzhou Top-Notch Talent Groups (ZXD2022003), the Shenzhen Medical Research Fund (B2403009), and the Sichuan Provincial Healthcare and Health Promotion Association (KY2024SJ0018).

## Author contributions

Z.Z., J.D., and M.W. conceptualized the project and designed the study. J.D., M.W., Y.J., H.D., R.Z., Y.L., and *Acinetobacter baumannii* Research Group contributed study materials. Y.Z., H.Lin, P.H., and M.W. performed the experiments. S.L., Y.W., Y.Z., L.Z., Y.W., J.L., H.Li, S.X., analyzed and interpreted data. S.L., Y.W., and Z.Z. visualized the results and prepared figures. S.L. wrote the initial draft of the manuscript. P.H., M.W., J.D., and Z.Z. discussed and revised the manuscript. All authors edited and commented on the manuscript.

## Ethics declaration

Animal experiments were approved by the Laboratory Animal Ethics Committee of Wenzhou Medical University (wydw2025-0449). Six-week old male Balb/c mice were obtained from Zhejiang Vital River Experimental Animal Technology Co., Ltd. (China) and maintained under specific pathogen-free (SPF) conditions. After one week of acclimatization, mice were randomly assigned into 7 groups (15 mice per group). For nasal colonization, 40 µL of the bacterial suspension was slowly administered intranasally (20 µL per nostril) to anesthetized mice to simulate upper respiratory tract colonization. Control mice received 40 µL PBS. The initial bacterial inoculum was quantified by serial dilution and plating.

Mice were euthanized and nasal tissues were collected at 1, 2, 3-, 5-, 7-, and 14-days post-inoculation respectively. Nasal mucosa tissues were aseptically excised and transferred into 1 mL PBS processed using a SCIENTZ-48 tissue grinder at 65 Hz, followed by immediate cooling on ice. Homogenate were serially diluted 10- fold with PBS. A total of 100 µL of homogenate or appropriate dilution was added to the surface of LB plate for colony counting, with 3 replicates for each sample.

## Supplementary Figures

**Fig. S1:**
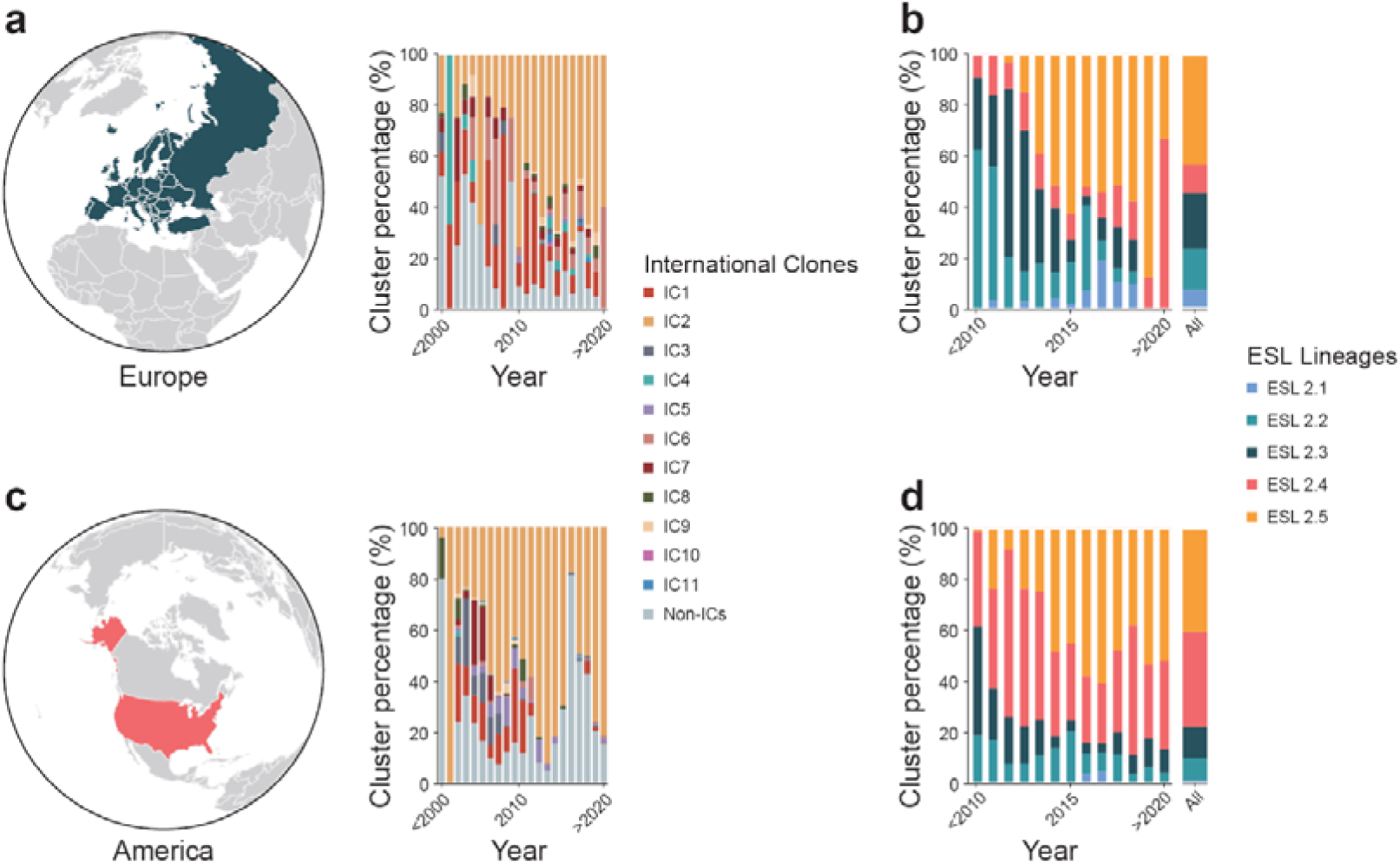
Temporal dynamics of international clones and ESL lineages in Europe and America. Geographic coverage and temporal distributions of ICs in Europe (**a**) and America (**c**). Temporal distributions of ESL lineages among IC2 isolates in Europe (**b**) and America (**d**).

**Fig. S2:**
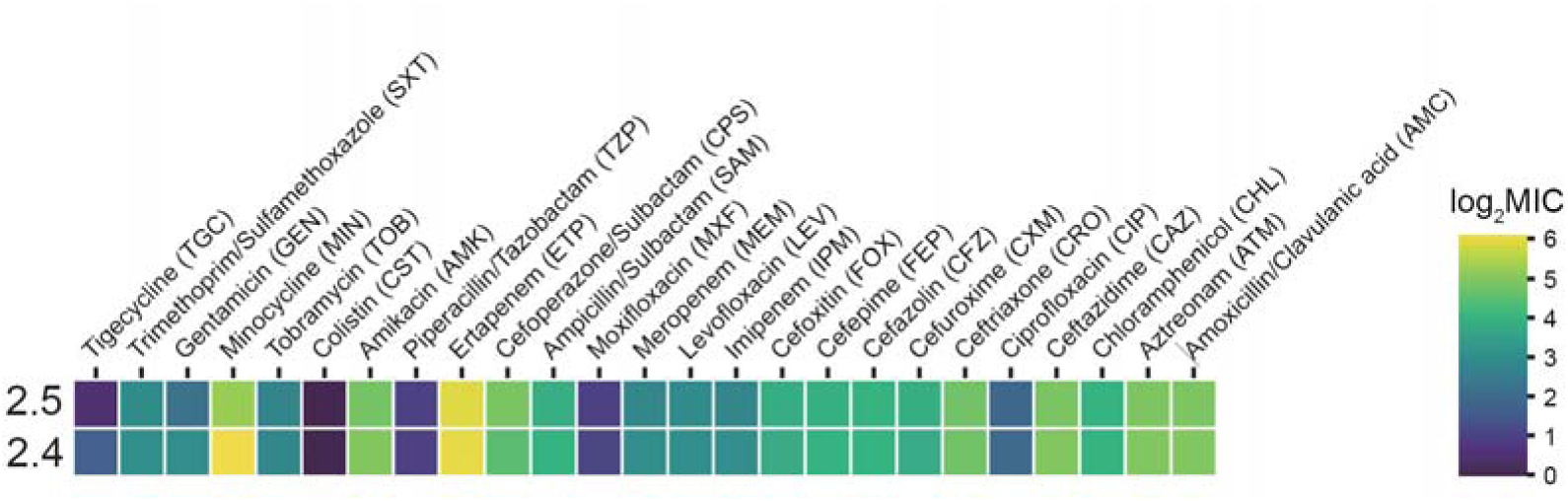
Antibiotic susceptibility profiles of ESL2.4 and ESL2.5 lineages. Heatmap showing the minimum inhibitory concentrations (MICs) of ESL2.4 and ESL2.5 isolates across a panel of clinically relevant antibiotics. Colours indicate log_-transformed MIC values.

**Fig. S3:**
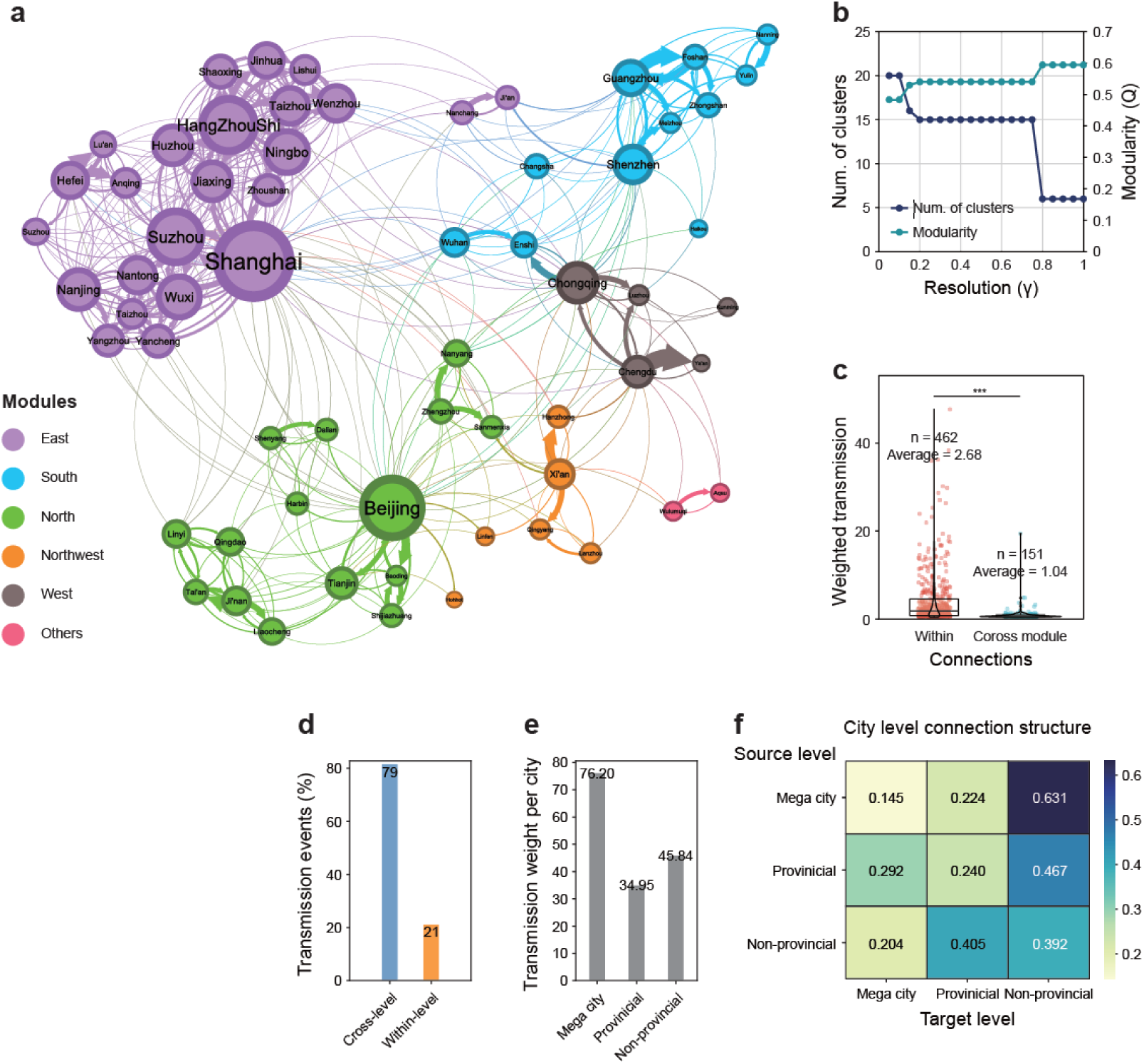
Modular structure and hierarchical transmission patterns of the city-level network. **a,** Module structure of the city-level transmission network. Nodes represent cities, edge weights indicate cumulative transmission events, and colors denote modules. **b,** Relationship between resolution parameter (γ) and the number of detected modules during community detection. **c,** Distribution of weighted transmission strength within modules and across modules. **d,** Comparison of transmission events occurring within city levels and across city levels. **e,** Average transmission weight per city stratified by city level. **f,** Proportional distribution of transmission events between source and target city levels.

**Fig. S4:**
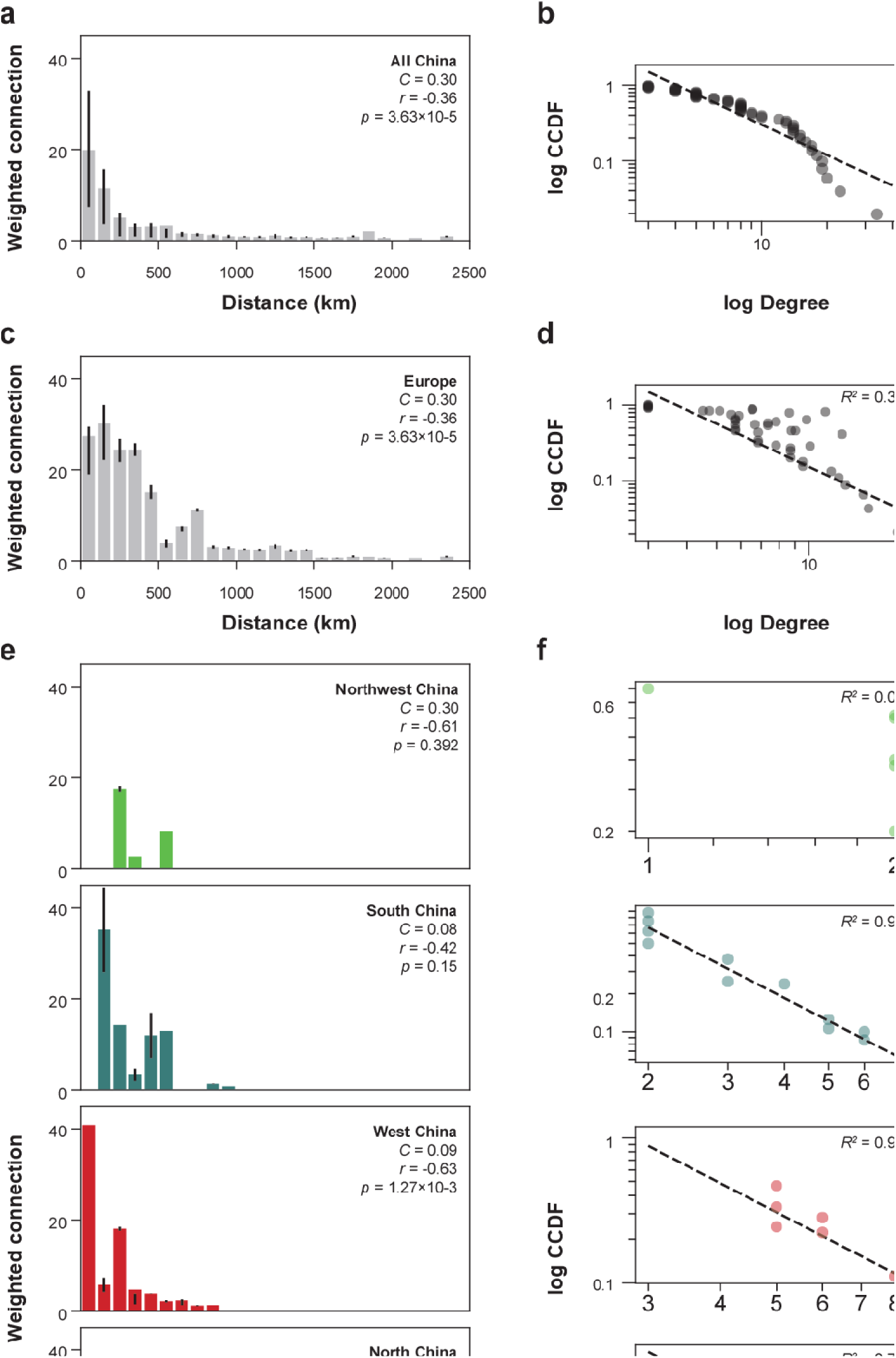
Distance-dependent transmission strength and degree distributions across regions. **a,c,e,** Histograms showing the relationship between geographic distance and weighted transmission strength for all of China (**a**), Europe (**c**), and major regions within China (**e**). Bars indicate mean weighted connections across distance bins. **b,d,f,** Power-law assessment of degree distributions corresponding to the networks shown in panels a, c, and e. Log–log complementary cumulative distribution functions (CCDFs) are shown with fitted trends and goodness-of-fit statistics.

**Fig. S5:**
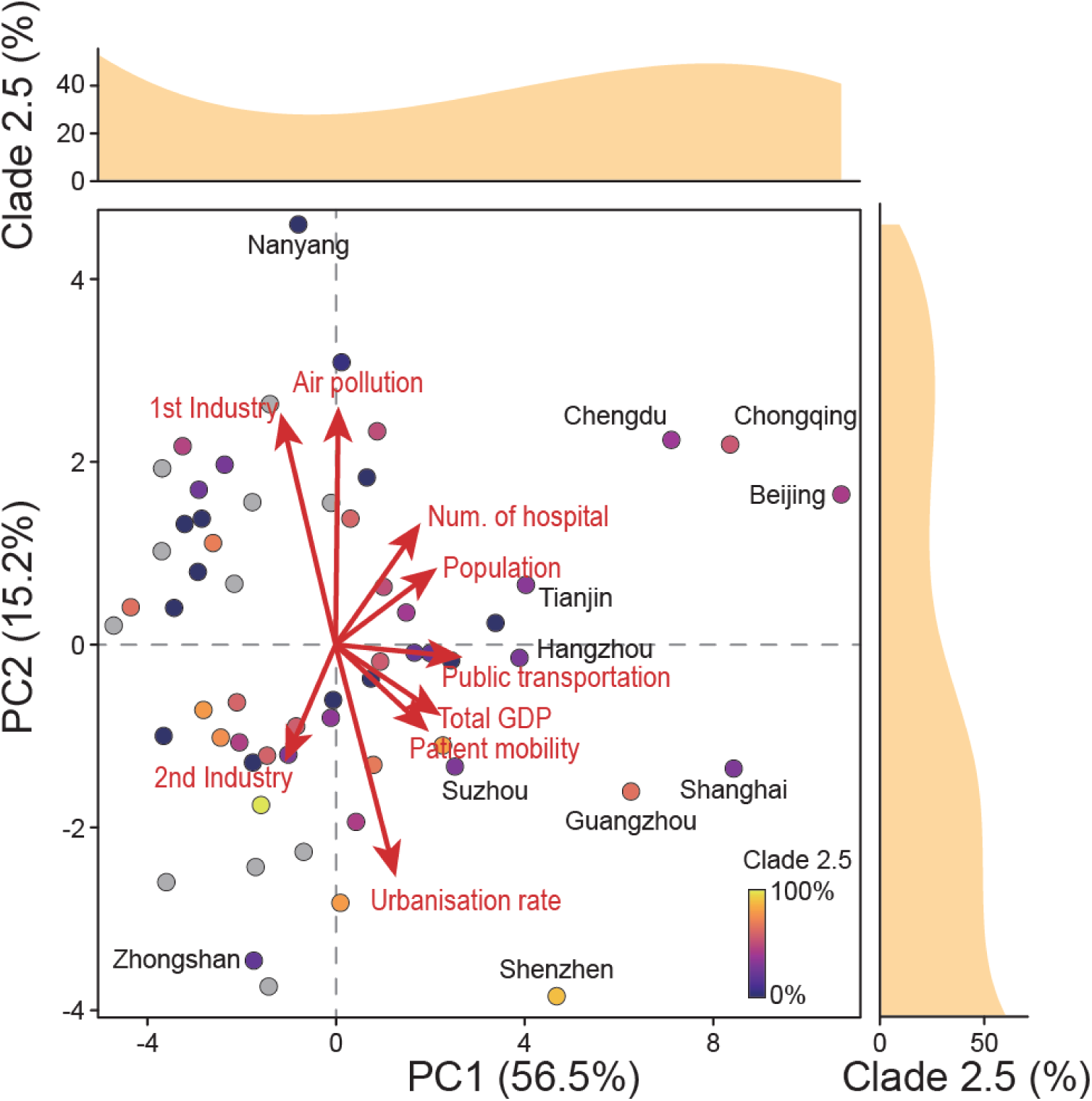
Redundancy analysis including all explanatory variables. Redundancy analysis (RDA) showing the full multivariable model with all variables displayed simultaneously. Arrows indicate the direction and relative contribution of each explanatory variable to the ordination, while points represent cities colored by the proportion of ESL2.5. Marginal density plots show the distribution of ESL2.5 prevalence along the first two RDA axes.

**Fig. S6:**
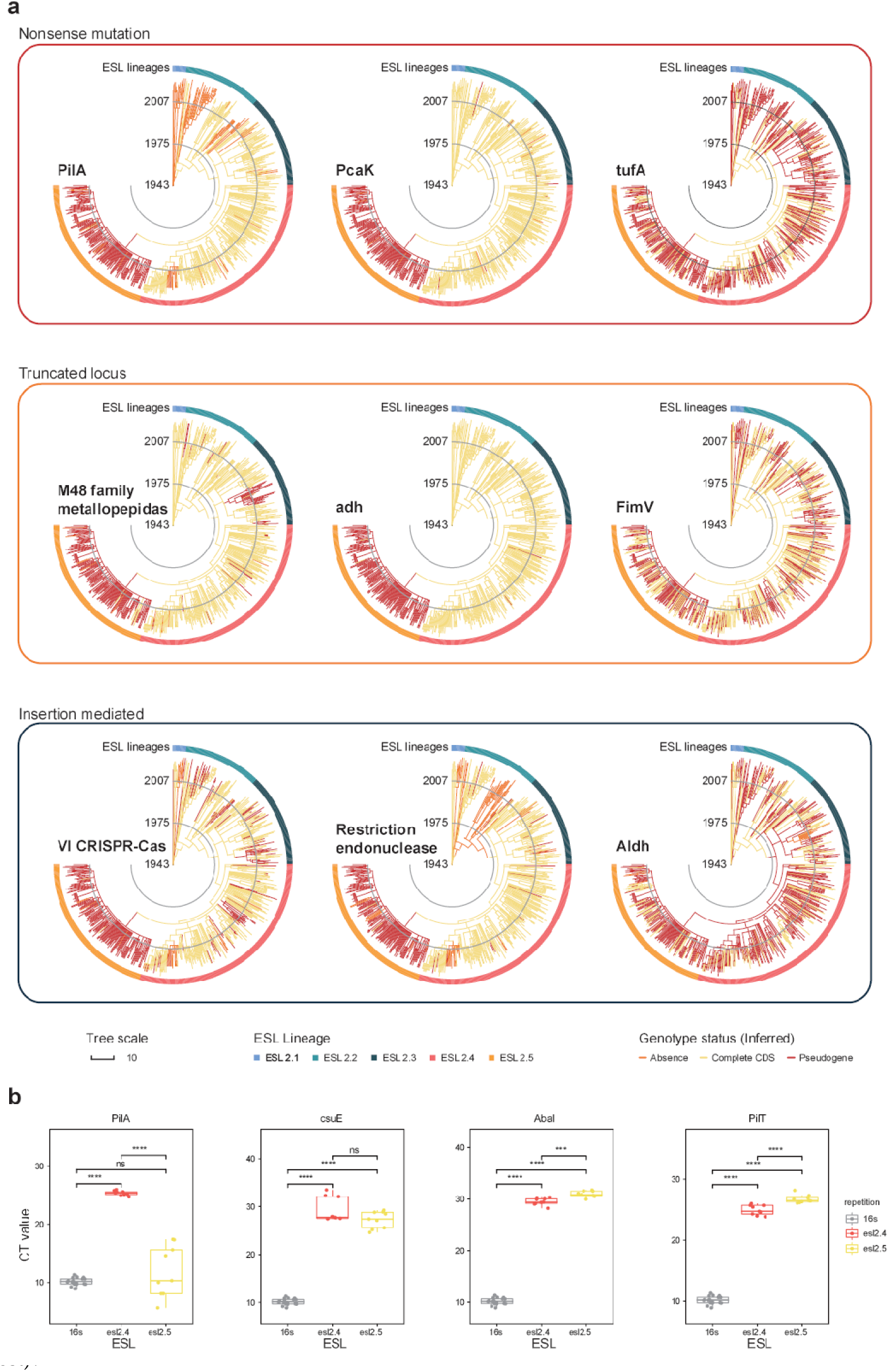
Evolutionary distribution and functional consequences of gene disruptions. **a,** Time-calibrated phylogenies inferred by TreeTime showing the evolutionary distribution of gene disruptions arising from three distinct mechanisms: nonsense mutations, locus truncation, and insertion-mediated disruption. Representative genes are shown for each category, and branches are colored by ESL lineage, while genotype status (absence, intact CDS, or pseudogene) is indicated along the outer ring. **b,** Quantitative PCR (qPCR) validation of selected disrupted genes across ESL lineages. Cycle threshold (CT) values are shown for intact and disrupted loci (\*\*\**P* < 0.001; \*\*\*\**P* < 0.0001; n.s. not significant; Mann-Whitney U test).

**Fig. S7:**
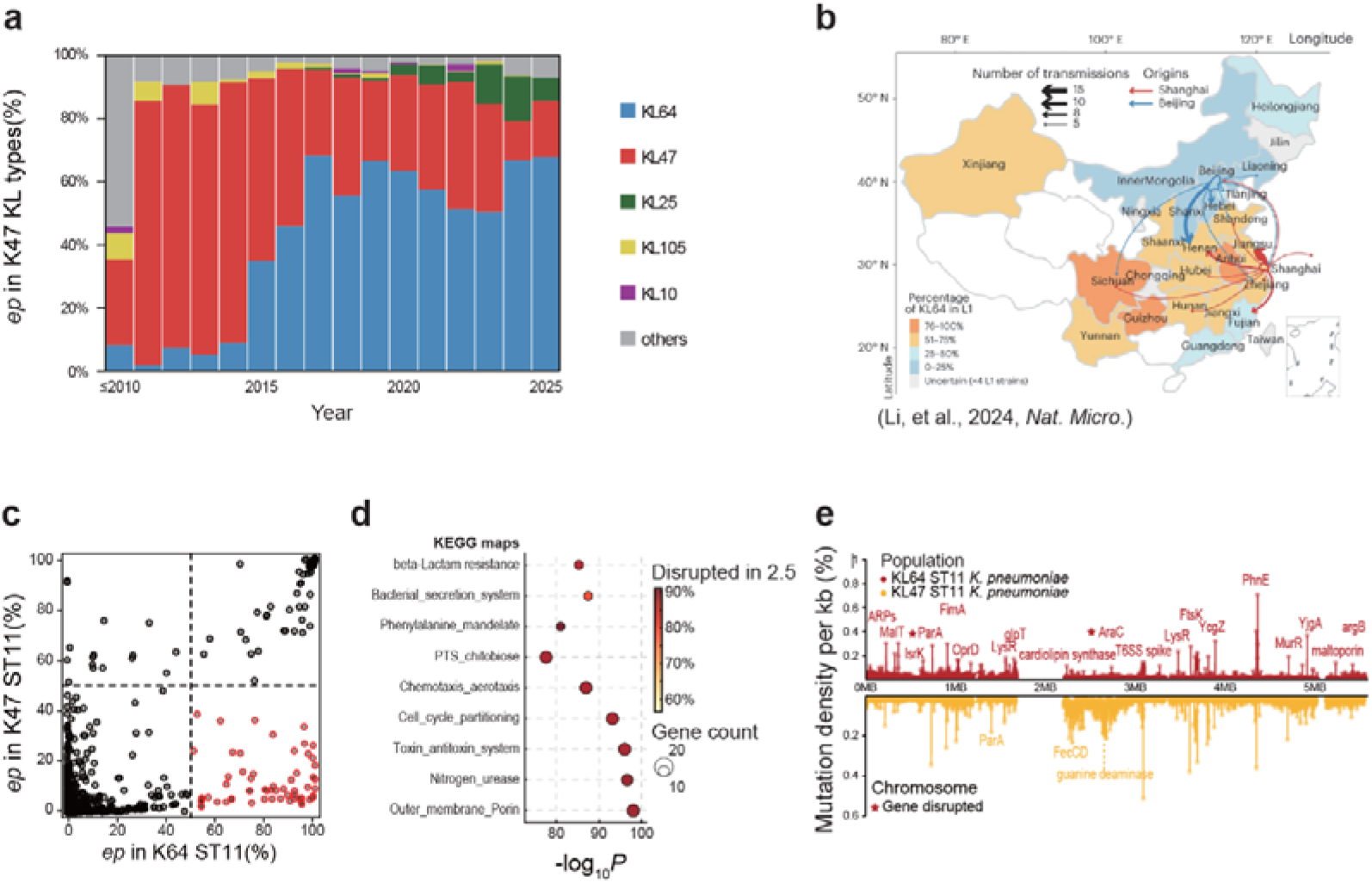
Parallel capsule dynamics, transmission pattern and genomic evolution in ST11 *K. pneumoniae* in China. **a,** Temporal changes in capsule locus composition within ST11 *K. pneumoniae*. **b,** Geographic distribution and inferred transmission routes of ST11 *K. pneumoniae* across China, with arrows indicating direction and relative frequency of interregional spread (adapted from Li *et al.*, 2024, *Nature Microbiology*). **c,** Gene-level disruption frequencies in KL64 ST11 *Klebsiella pneumoniae* relative to non-KL64 lineages. Each point represents a gene, plotted by the proportion of disrupted coding sequences in KL64 (x-axis) versus non-KL64 ST11 lineages (y-axis). Genes significantly enriched for disruption in KL64 are highlighted. **d,** Functional enrichment analysis of genes preferentially disrupted in the KL64 ST11 lineage. **e,** Genome-wide distribution of mutation density in ST11 *K. pneumoniae*, with mutation hotspots and representative disrupted genes indicated along the chromosome.

