## Supplementary Figures for "Healthcare Infrastructure Shapes Evolutionary Trade-offs and Geographic Dissemination of Multidrug-Resistant *Acinetobacter baumannii*"

Iotabiome Biotechnology Inc: Shichang Xie, Yi Ren

West China Second University Hospital: Linghan Kuang

The First Affiliated Hospital of Hebei North University: Li Wang

Heilongjiang General Hospital of Daqing Oil Field: Min Liu, Jinwen Wang

Weifang People's Hospital: Wanxiang Li

No. 3201 Hospital: Yihai Gu, Wei Zhang

Jiaxing TCM Hospital Affiliated to Zhejiang Chinese Medical University: Xingying Chen

Taizhou Hospital: Sufei Yu

The First Hospital of Ningbo: Wei Liang

The Second Hospital of Yinzhou, Ningbo: Shengke Wang

The Second Hospital of Shaoxing: Chaochao Wang

Wenzhou Central Hospital: Yangfang Chen

The Third People’s Hospital of Tianjin: Sumei Wang

Tianjin First Central Hospital: Jingyu Wang

Shenzhen Center for Disease Control and Prevention: Yinghui Li, Miaoling Chen, Lulu Hu

**Supplementary Figures**

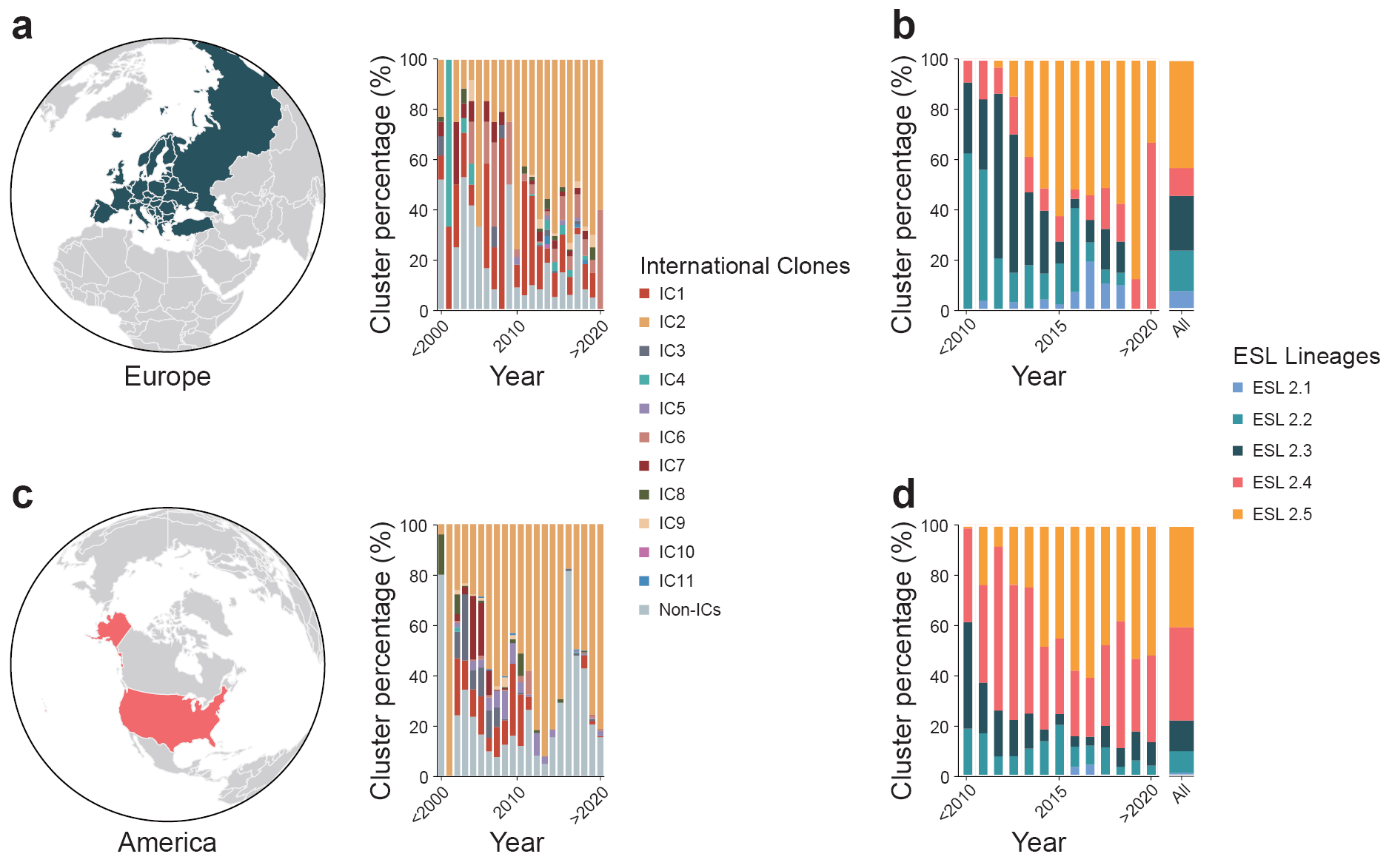

**Fig. S1: Temporal dynamics of international clones and ESL lineages in Europe and America**

Geographic coverage and temporal distributions of ICs in Europe (**a**) and America (**c**). Temporal distributions of ESL lineages among IC2 isolates in Europe (**b**) and America (**d**).

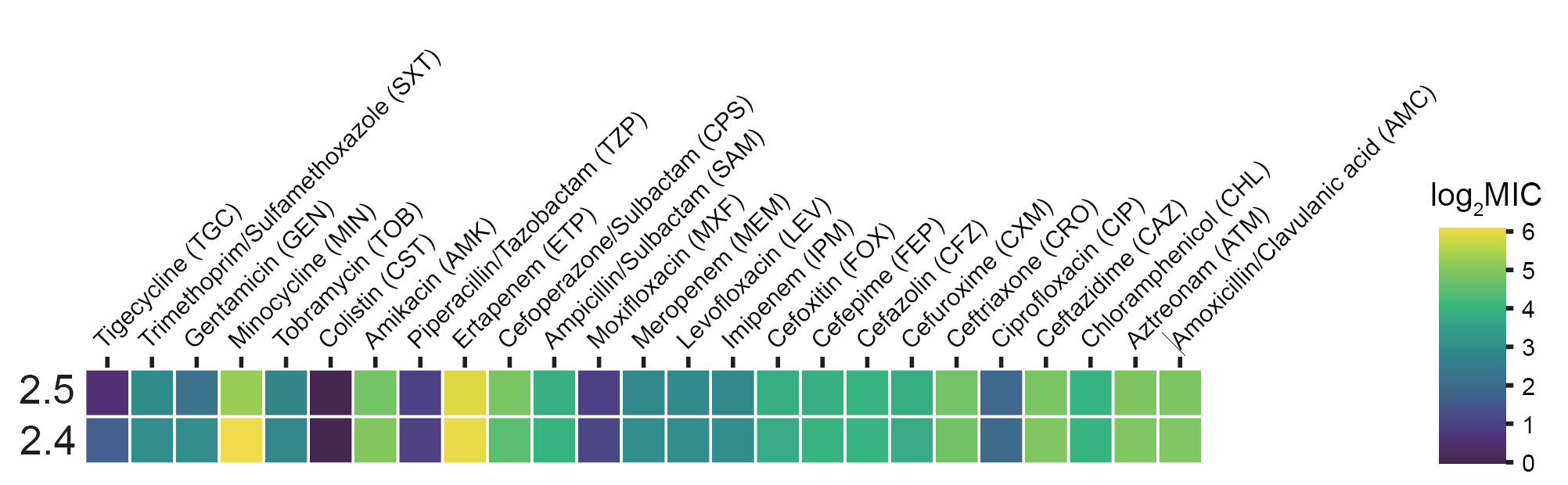

**Fig. S2: Antibiotic susceptibility profiles of ESL2.4 and ESL2.5 lineages**

Heatmap showing the minimum inhibitory concentrations (MICs) of ESL2.4 and ESL2.5 isolates across a panel of clinically relevant antibiotics. Colours indicate log₂-transformed MIC values.

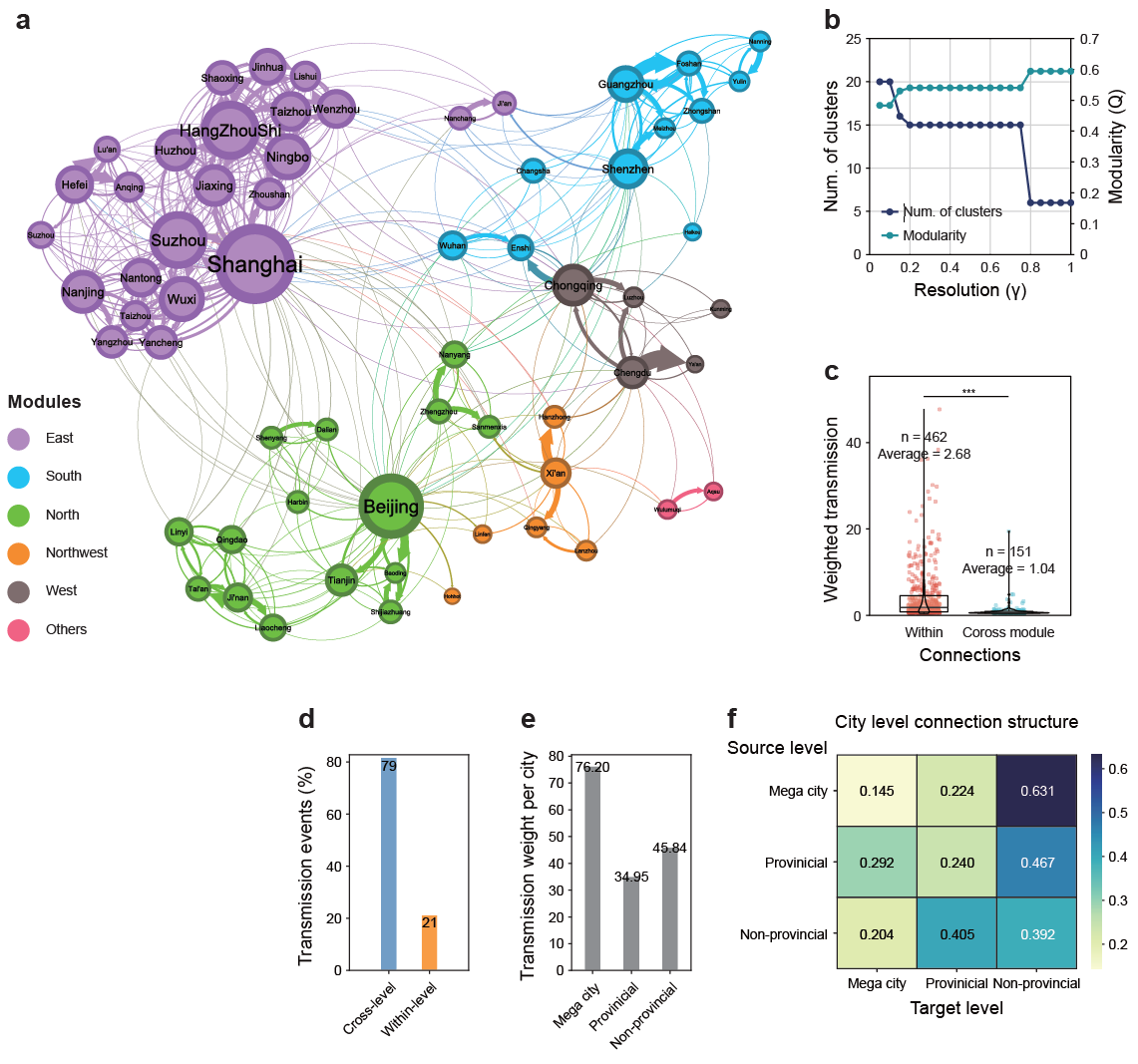

**Fig. S3: Modular structure and hierarchical transmission patterns of the city-level network**

**a,** Module structure of the city-level transmission network. Nodes represent cities, edge weights indicate cumulative transmission events, and colors denote modules. **b,** Relationship between resolution parameter (γ) and the number of detected modules during community detection. **c,** Distribution of weighted transmission strength within modules and across modules. **d,** Comparison of transmission events occurring within city levels and across city levels. **e,** Average transmission weight per city stratified by city level. **f,** Proportional distribution of transmission events between source and target city levels.

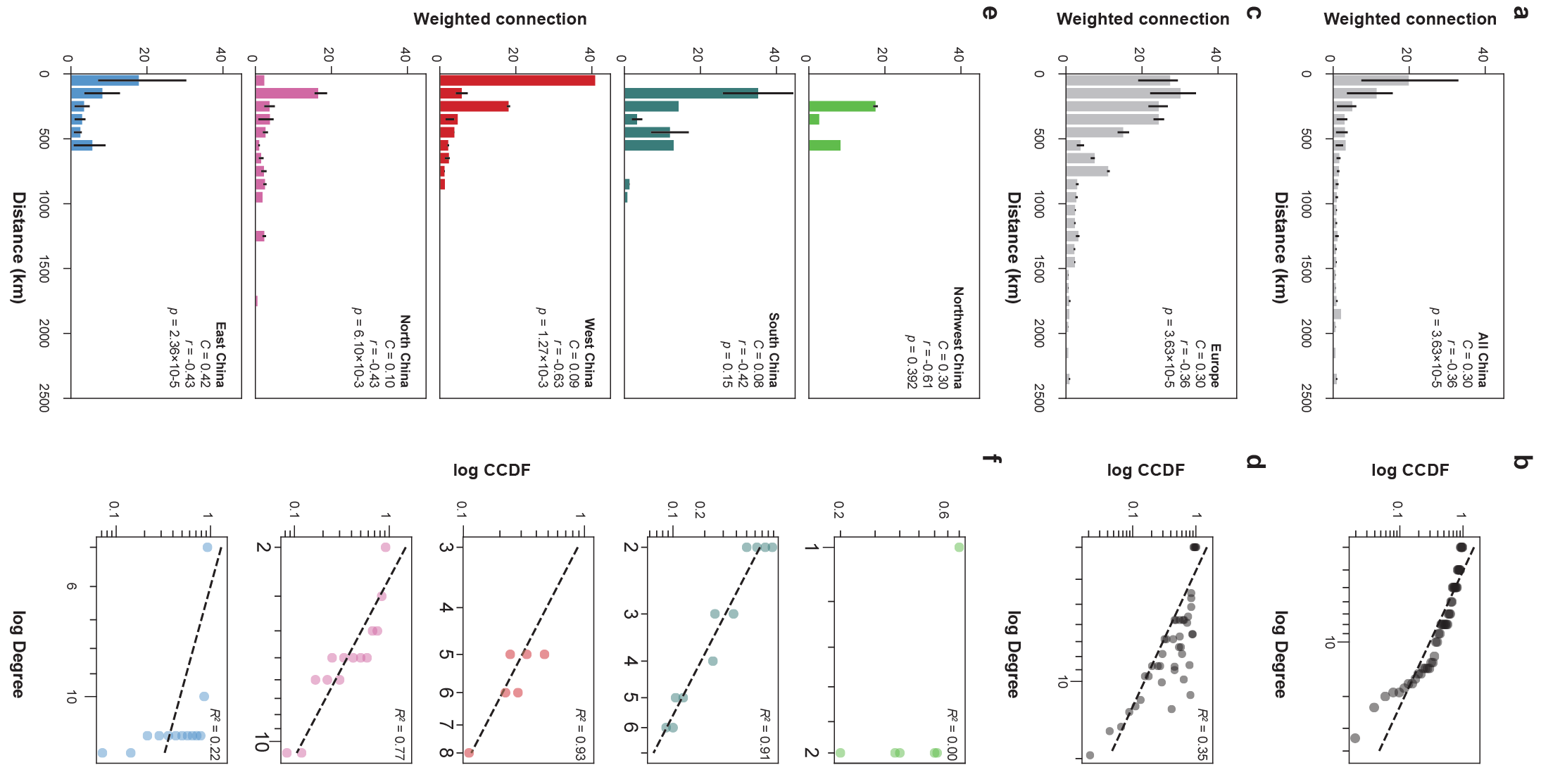

**Fig. S4: Distance-dependent transmission strength and degree distributions across regions**

**a,c,e,** Histograms showing the relationship between geographic distance and weighted transmission strength for all of China (**a**), Europe (**c**), and major regions within China (**e**). Bars indicate mean weighted connections across distance bins. **b,d,f,** Power-law assessment of degree distributions corresponding to the networks shown in panels a, c, and e. Log–log complementary cumulative distribution functions (CCDFs) are shown with fitted trends and goodness-of-fit statistics.

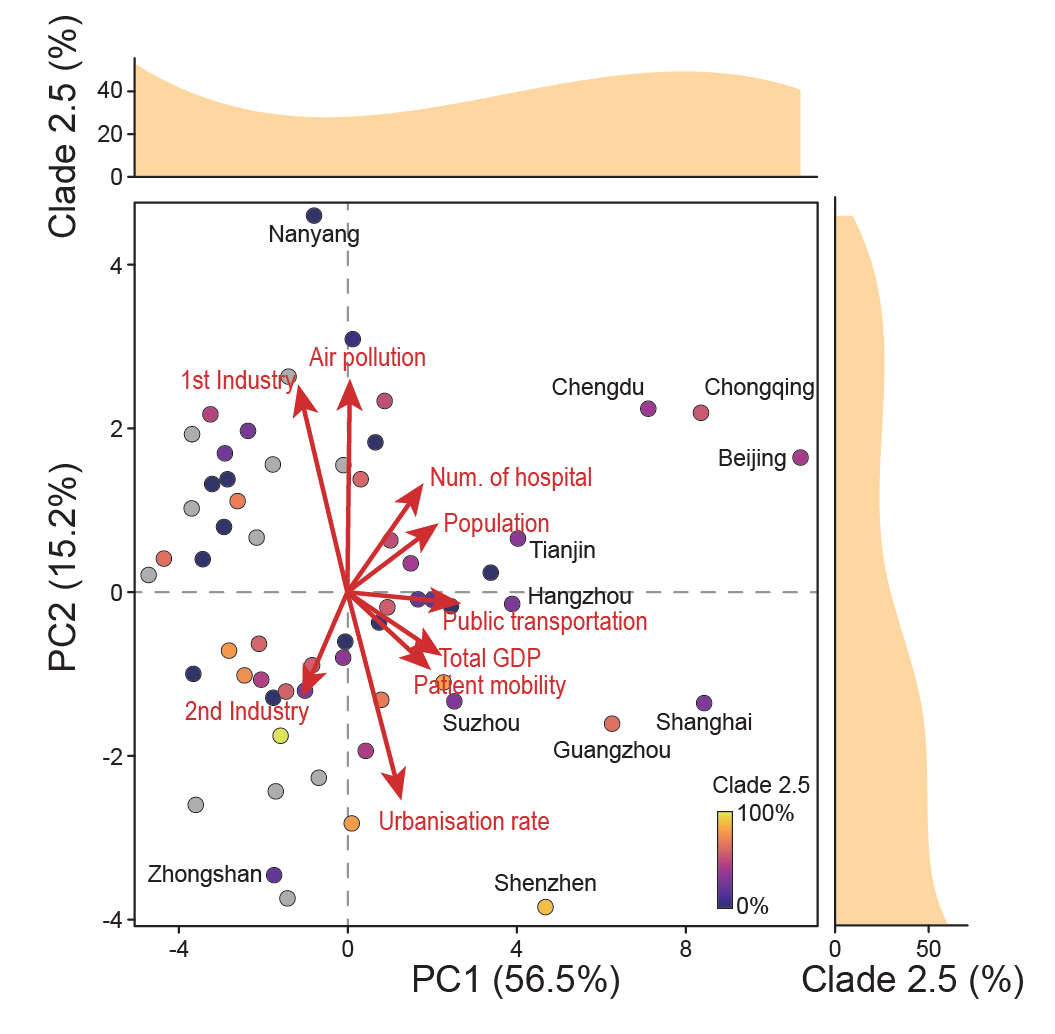

**Fig. S5: Redundancy analysis including all explanatory variables**

Redundancy analysis (RDA) showing the full multivariable model with all variables displayed simultaneously. Arrows indicate the direction and relative contribution of each explanatory variable to the ordination, while points represent cities colored by the proportion of ESL2.5. Marginal density plots show the distribution of ESL2.5 prevalence along the first two RDA axes.

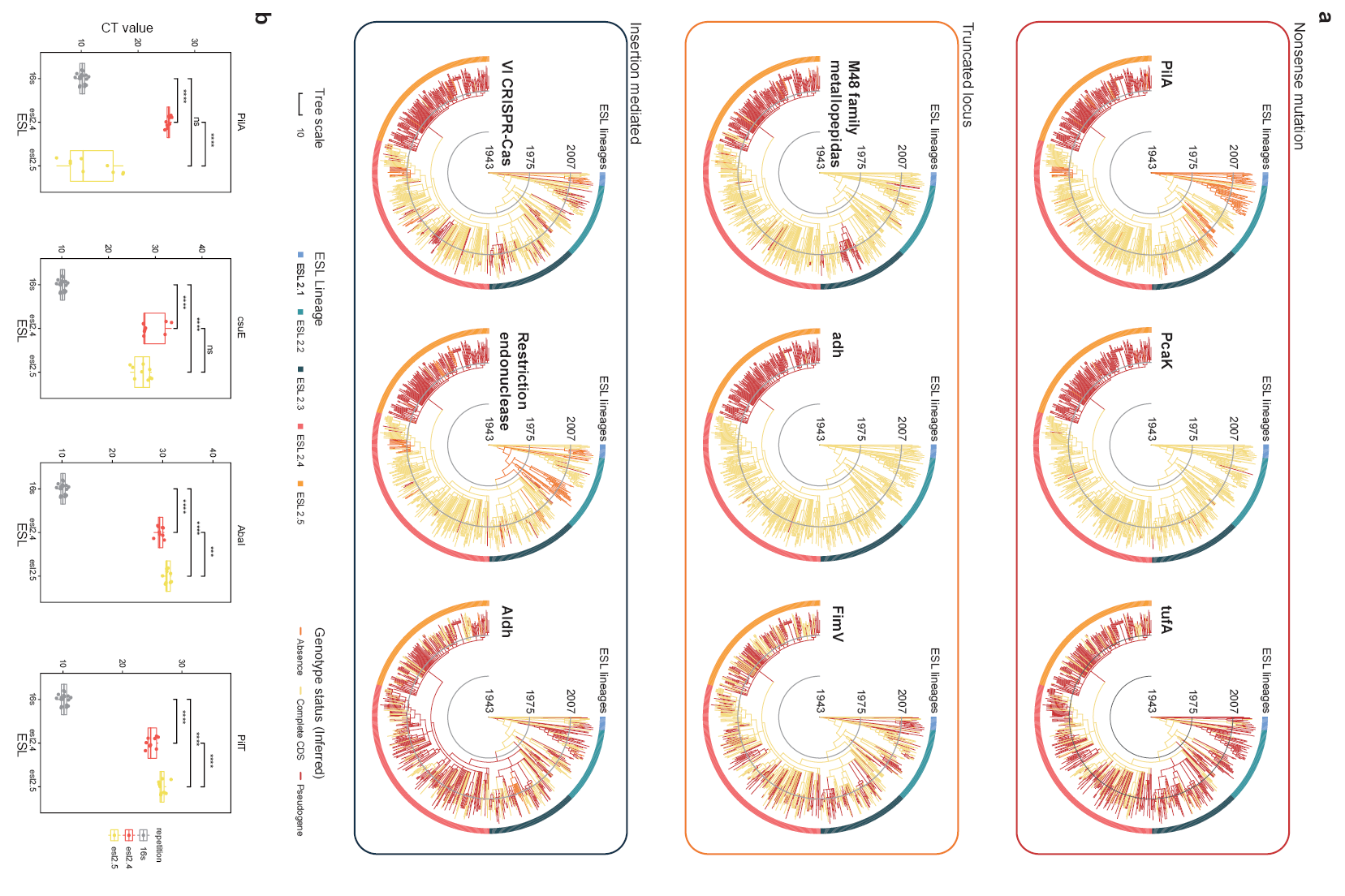

**Fig. S6: Evolutionary distribution and functional consequences of gene disruptions**

**a,** Time-calibrated phylogenies inferred by TreeTime showing the evolutionary distribution of gene disruptions arising from three distinct mechanisms: nonsense mutations, locus truncation, and insertion-mediated disruption. Representative genes are shown for each category, and branches are colored by ESL lineage, while genotype status (absence, intact CDS, or pseudogene) is indicated along the outer ring. **b,** Quantitative PCR (qPCR) validation of selected disrupted genes across ESL lineages. Cycle threshold (CT) values are shown for intact and disrupted loci (****P* < 0.001; *****P* < 0.0001; n.s. not significant; Mann-Whitney U test).

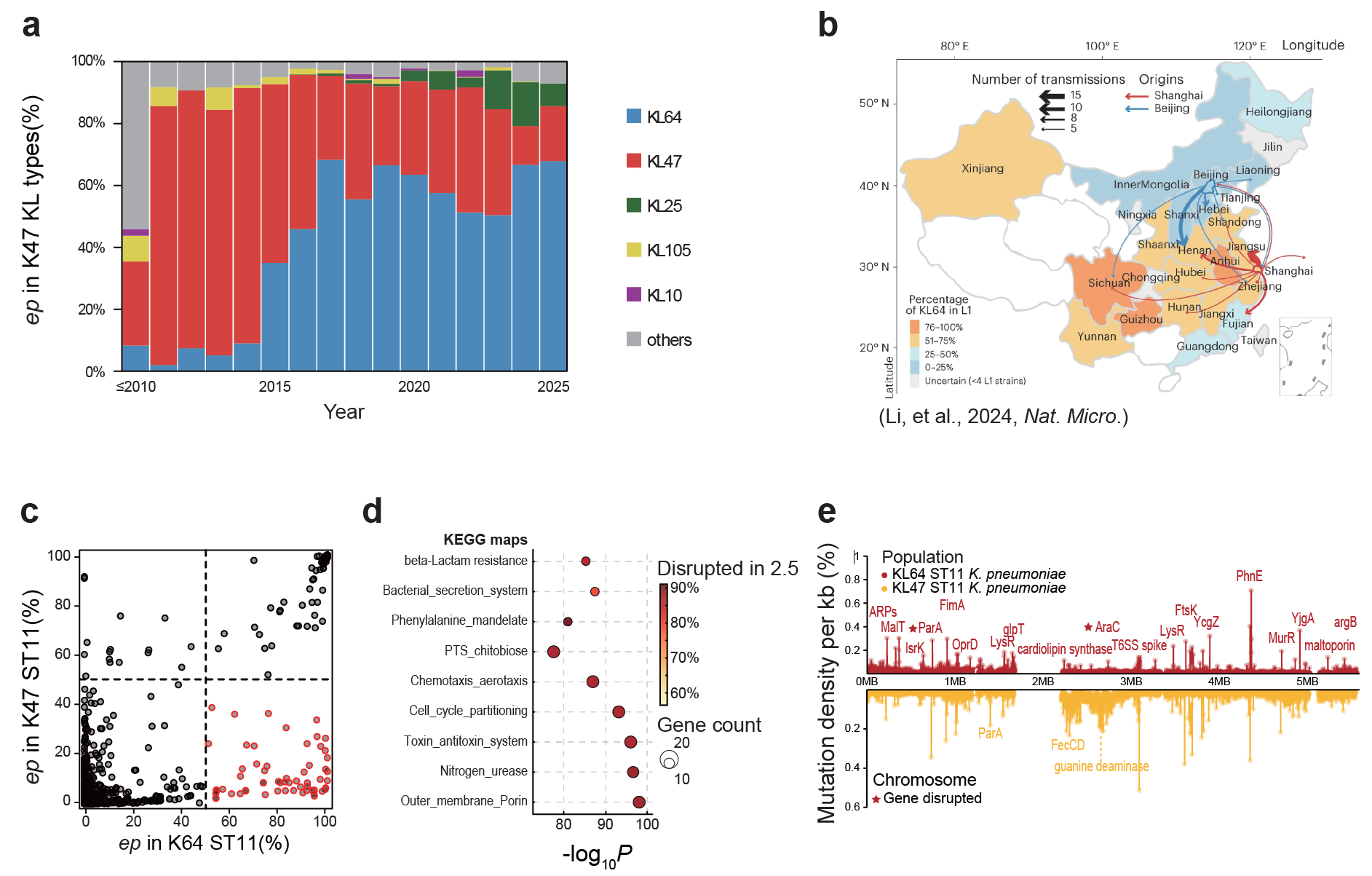

**Fig. S7: Parallel capsule dynamics, transmission pattern and genomic evolution in ST11 *K. pneumoniae* in China**

**a,** Temporal changes in capsule locus composition within ST11 *K. pneumoniae*. **b,** Geographic distribution and inferred transmission routes of ST11 *K. pneumoniae* across China, with arrows indicating direction and relative frequency of interregional spread (adapted from Li *et al.*, 2024, *Nature Microbiology*). **c,** Gene-level disruption frequencies in KL64 ST11 *Klebsiella pneumoniae* relative to non-KL64 lineages. Each point represents a gene, plotted by the proportion of disrupted coding sequences in KL64 (x-axis) versus non-KL64 ST11 lineages (y-axis). Genes significantly enriched for disruption in KL64 are highlighted. **d,** Functional enrichment analysis of genes preferentially disrupted in the KL64 ST11 lineage. **e,** Genome-wide distribution of mutation density in ST11 *K. pneumoniae*, with mutation hotspots and representative disrupted genes indicated along the chromosome.
